# Trajectory-informed gene feature selection in single-cell analysis with SEEK-VFI

**DOI:** 10.64898/2025.12.12.694028

**Authors:** Rebecca Danning, Zheng Tracy Ke, Xihong Lin, Rong Ma

## Abstract

The prioritization of highly-variable genes is an important step in single-cell trajectory inference. However, when variability arises from a continuous latent cell development trajectory, standard methods may fail to differentiate trajectory-relevant from uninformative genes. SEEK-VFI is an ensemble topic-modeling machine learning algorithm for trajectory inference preprocessing that prioritizes trajectory-relevant genes. It outperforms existing methods, and identifies key genes that improve trajectory topology reconstruction, enhance visualization, and augment downstream trajectory analyses.

---

Single-cell trajectory inference aims to understand the developmental structure underlying cell differentiation based on cell-level gene expression profiles [1]. Trajectory inference methods typically represent the trajectory as a graph comprising developmental milestones (nodes) and transitions (edges) between adjacent milestones [2]. Feature selection is a critical preprocessing step in trajectory inference, and is typically done by choosing genes with the greatest variability in expression across cells [3]. A fundamental assumption of trajectory inference is that cells lie along continuous, latent transitions among cellular states that give rise to variation in gene expression [4]. However, standard feature selection methods [5-7] do not exploit the underlying low-dimensional, smooth structure of the expression data; instead, they operate directly on the high-dimensional normalized expression matrices. These approaches are primarily designed to select genes that are differentially expressed across discrete cell clusters, such as distinct cell types [3, 8]. Consequently, when applied to trajectory inference, existing feature selection methods often fail to capture genes that play key roles in developmental dynamics, particularly those whose expression changes gradually along continuous differentiation trajectories rather than sharply between clusters.

To address this limitation, here we introduce SEEK-VFI (Spectral Ensembling of topic models with Eigenscore for K-agnostic Variable Feature Identification), an ensemble machine learning method for prioritizing genes that are highly variable with respect to latent topological structure. Building on topic modeling, SEEK-VFI enables efficient detection of trajectory-associated features through its flexible representation of mixed cellular memberships along continuous developmental trajectories. SEEK-VFI is a preprocessing method and thus does not assume the availability of a cell embedding representing the complete trajectory; rather, it leverages latent trajectory topology indirectly to inform feature prioritization. An overview of topic modeling for trajectory inference and the SEEK-VFI workflow is shown in Fig. 1.

**Fig 1.**
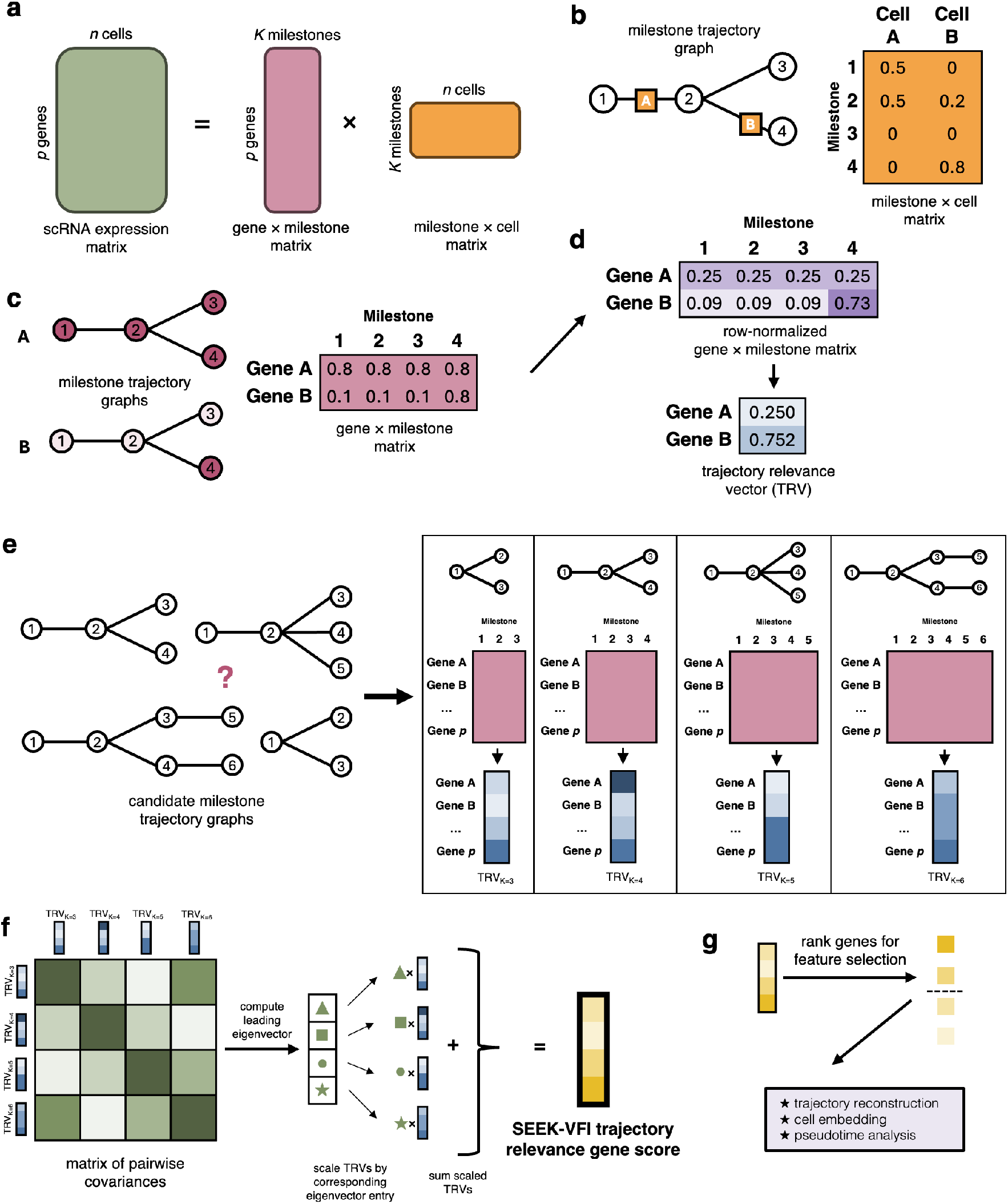
SEEK-VFI workflow for prioritizing trajectory-relevant genes. a)Topic modeling decomposes the gene × cell expression matrix into two low-rank matrices defined by the latent trajectory structure. b) The milestone × cell matrix describes the position of each cell with respect to its trajectory segment. In the diagram, Cell A lies halfway between milestones 1 and 2, while Cell B lies closer to milestone 4 than 2. c) The gene × milestone matrix describes gene expression patterns with respect to location on the trajectory. Gene A has high expression across the entire trajectory, while Gene B has high expression at milestone 4 and low expression otherwise. d) The gene × milestone matrix can be normalized by dividing each row by its *ℓ*^1^-norm. The normalized row corresponding to Gene A is uniform, while the normalized row corresponding to Gene B is spikier; Gene B is therefore the gene that is more relevant to trajectory structure. A candidate trajectory relevance score measuring spikiness is calculated for each gene by taking the squared *ℓ*^2^-norm of each normalized row (see Methods). e) Topic modeling requires the dimension of the latent trajectory space *K*. Since the underlying trajectory structure is unknown, we require a method that ensembles the spikiness vectors from a variety of candidate trajectories. f) SEEK-VFI combines the candidate trajectory relevance vectors from a range of topic models using a spectral ensembling method that yields a robust and trajectory-agnostic estimator of gene relevance. g) Feature selection with SEEK-VFI enables downstream tasks such as trajectory reconstruction, cell embedding, and pseudotime analysis.

Topic modeling assumes a low-rank structure underlying the expression matrix [9] and seeks to decompose the gene × cell matrix into a gene × milestone and milestone × cell matrix (Fig. 1a). The milestone × cell matrix captures the position of each cell along the trajectory (Fig. 1b), whereas the gene × milestone matrix captures the expected expression level of each gene at each milestone (Fig. 1c). Intuitively, if Gene A shows relatively uniform expression across all milestones along the trajectory, while gene B is highly expressed only at a specific milestone, the variation in gene B’s expression across cells (“spikiness”) would be more biologically meaningful and informative of the underlying developmental process (Fig. 1c).

To quantify the biological relevance of each gene with respect to the underlying trajectory, we compute the spikiness of each gene from the estimated gene × milestone matrix. The spikiness metric is designed to highlight genes whose expression profiles across milestones contain one or two dominant loadings and much smaller values in the remaining components (Methods). Genes with high spikiness vary more significantly along the trajectory and are prioritized for downstream analysis, while genes with a low spikiness exhibit relatively uniform expression along the trajectory and are less informative. These measures are collected into a trajectory relevance vector (TRV) (Fig. 1d).

SEEK-VFI uses Topic-SCORE [9], an efficient SVD-based method, to derive the gene × milestone matrices. Topic modeling algorithms require the specification of *K*, the number of milestones; however, this is often unknown. Thus, the first step of SEEK-VFI is to run Topic-SCORE corresponding to a range of plausible values of *K* (Fig. 1e). For each candidate value of *K*, we can compute a TRV that summarizes each gene’s importance with respect to that milestone model. These vectors are then aggregated using an optimal spectral ensemble method [10, 11] (Fig. 1f). The output of SEEK-VFI is a consensus trajectory relevance score for each gene. These scores can be used to rank the genes and guide feature selection in downstream analysis (Fig. 1g).

To benchmark SEEK-VFI’s ability to distinguish trajectory-relevant genes from uninformative ones, we first test it on simulated trajectory data and compare it to the three existing methods available in Seurat and the recent method DELVE [12]. The simulated data are generated such that 10% of the genes have expression levels related to the milestone structure (trajectory-relevant) and 90% of genes have constant expression levels across the trajectory (uninformative). We vary the signal strength parameter *α*, which dictates how much the expression levels of the trajectory-relevant genes vary across milestones; further details are provided in Methods. Fig. 2a shows the AUC for SEEK-VFI and the alternative methods in identifying the trajectory-relevant genes across the settings. We found SEEK-VFI consistently outperformed the alternative methods, particularly with increased sample size (Supplemental Fig. S3). While DELVE performs comparably to the Seurat methods, analysis of a single dataset containing only thousands of cells takes hours, so we chose to exclude it from further analyses. Notably, SEEK-VFI is the most robust to different trajectory topological structures, which can be seen clearly in Supplemental Fig. S4.

**Fig. 2.**
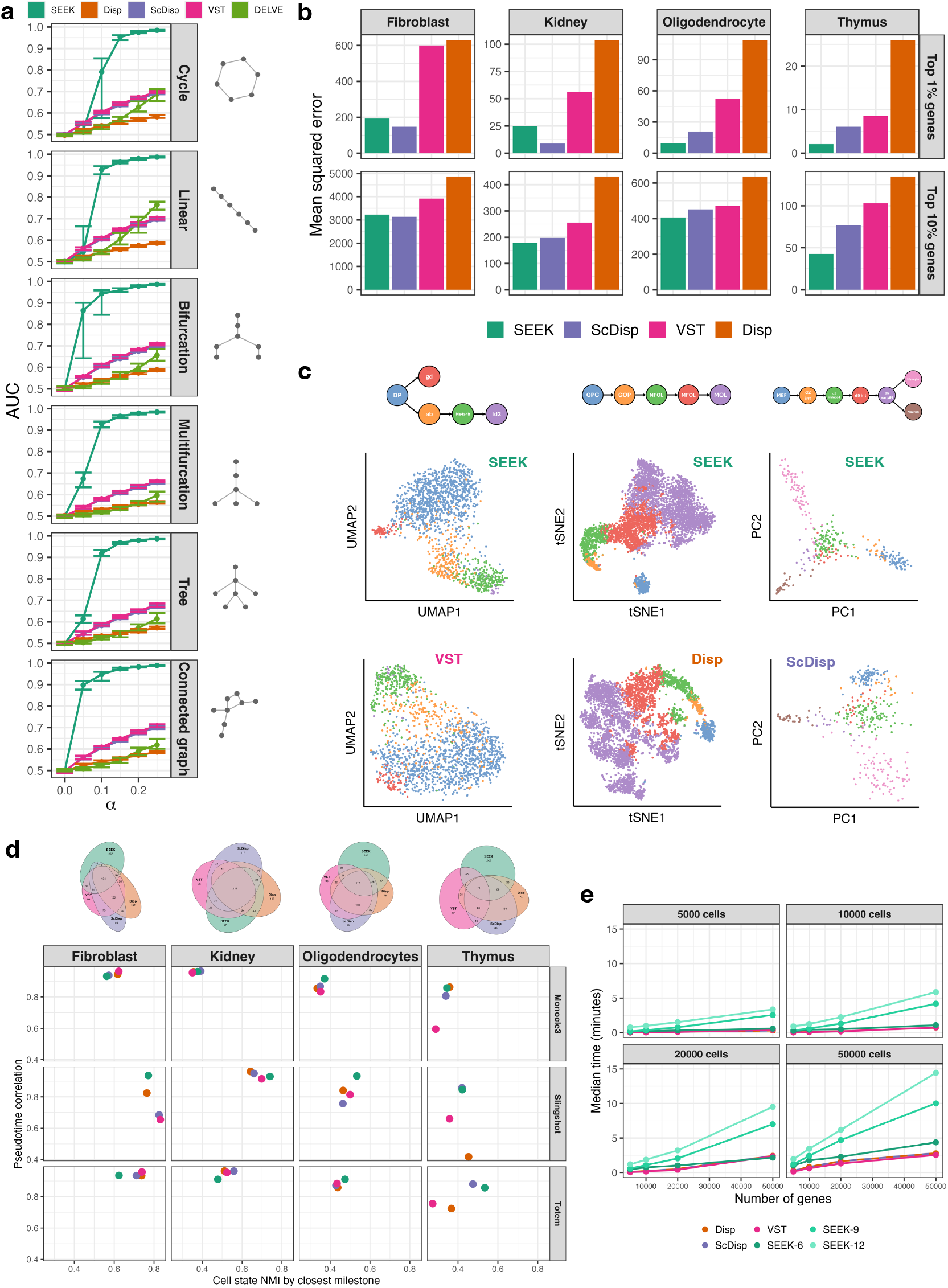
Comparison with existing methods in simulated and real datasets. a) Performance of SEEK-VFI and alternative methods on simulated scRNA-seq data with 5,000 cells and 10,000 genes. The x-axis varies the value of *α*, which controls how similar the trajectory-relevant genes are to the uninformative genes. The y-axis shows the median AUC for differentiating between informative and uninformative genes across the simulations. Error bars represent the 25^*th*^ and 75^*th*^ percentiles. Results from expanded settings are shown in Supplemental Fig. S3 and Supplemental Fig. S4. b) Bar plots showing the mean squared error comparing the true cell-to-cell trajectory distance matrices (see Methods) to the distance matrices derived from the normalized expression of the top X% genes as selected by each of the four methods. Results for additional datasets and values of X are shown in Supplemental Fig. S5. c) Low-dimensional embeddings using the top 500 genes as selected by the given method. The top row shows the ground-truth trajectory. Left: bifurcation of t-cells in the thymus, comparing the UMAP embeddings from SEEK-VFI and VST. Middle: linear trajectory of oligodencrocytes, comparing the t-SNE embeddings from SEEK-VFI and Disp. Right: reprogramming of fibroblast cells, comparing the PCA embeddings from SEEK-VFI and ScDisp. Results from an expanded set of visualization settings are found in Supplemental Figs S7-S30. d) Performance of downstream trajectory analysis tools using the top 500 genes from each method and Venn diagrams showing the overlap of the gene sets. e) Runtime of each method. SEEK-X ensembles the spikiness vectors from topic models with *K* = {3, …, *X*} milestones. Run-times for Seurat methods do not include normalization preprocessing. DELVE is omitted from panels (b)–(d) due to its long running time.

We test these methods on five real scRNA-seq datasets covering diverse developmental processes, comparing their performance in prioritizing the genes which best capture the underlying true trajectory (Fig. 2b). Across all datasets and various gene selection cutoffs, SEEK-VFI robustly achieves the overall lowest reconstruction error of the latent trajectory. This superiority is further reflected in the various low-dimensional embeddings of the cells based on the top 500 genes selected by each method (Fig. 2c).

We also demonstrate the advantages of SEEK-VFI over the existing methods in improving downstream tasks such as pseudotime inference and cell-state identification. For each of three popular trajectory inference methods (Monocle3 [13], Slingshot [14], and Totem [15]), we restrict the input to the expression profiles of the top 500 genes selected by each feature selection method. We then evaluate (1) the correlation of the derived pseudotime to the ground truth pseudotime, and (2) the concordance between the inferred milestone structure and the true cell states (see Methods). As shown in Fig. 2d, SEEK-VFI achieved the best overall performance across all datasets and trajectory inference algorithms. It performed comparably to existing feature selection methods on datasets with already strong baseline performance (e.g., kidney, hematopoiesis), while showing substantial improvement on more challenging datasets (e.g., fibroblast, thymus). Notably, there is smaller overlap between the genes prioritized by SEEK-VFI and the existing methods, as compared with the overlap among the existing methods themselves, suggesting the unique strength of SEEK-VFI in detecting distinct type of expression patterns commonly overlooked by conventional approaches (Fig. 2d).

The runtime of SEEK-VFI depends on dataset size and the number of candidate milestone models included in the ensemble (Fig. 2e). Although the conventional methods that do not require model estimation are fastest, the additional computational cost of SEEK-VFI is modest. For comparison, the other trajectory-aware algorithm DELVE took nearly 3 hours to analyze datasets with 5K cells and 10K genes. In our experiments, even for datasets containing 50K cells and over 20K genes, with as many as 10 candidate models and up to 12 milestones, SEEK-VFI completes within minutes on a standard laptop (MacBook Pro, 6-core Intel i7).

In sum, SEEK-VFI is a novel and powerful method for detecting trajectory-associated genes from single-cell transcriptomics data. We show through simulations that SEEK-VFI outperforms conventional feature selection methods at differentiating between trajectory-relevant and uninformative genes across a range of dataset sizes and cell trajectory structures. SEEK-VFI prioritizes trajectory-relevant genes in real scRNA-seq datasets across a variety of biological processes and trajectory structures, improving both trajectory embedding and pseudotime estimation. While SEEK-VFI shows promise for improving trajectory inference, there are some limitations to consider. First, the computational cost of the underlying Topic-SCORE algorithm increases exponentially with *K* (Supplemental Fig. S2a). Second, the underlying Topic-SCORE algorithm utilizes kmeans, which may occasionally suffer from instability in convergence. Third, the researcher must independently determine the range of candidate values of *K*. Finally, as with traditional feature selection methods, SEEK-VFI does not provide a guaranteed cutoff for the number of genes to select.

## Methods

Trajectory inference takes as its input a matrix **X** ∈ ℝ^*pxn*^ of *n* cells and *p* genes, where **X**_*ij*_ is cell *j*’s expression of gene *i* at a particular point in time. In the following section, we will describe SEEK-VFI in more detail, including how we apply the topic modeling framework to the trajectory inference context.

### Topic modeling and Topic-SCORE

Topic modeling is a statistical method from natural language processing that aims to find low-rank latent structure (“topics”) that explain the counts of vocabulary words in a collection of documents: vocabulary words have varying likelihoods of appearing across topics, and documents comprise a mixture of topics [16]. The observed matrix **X** ∈ ℝ^*pxn*^ summarizes the counts of *p* vocabulary words across *n* documents. Given a document *i* containing *n*_*i*_ words, the *i*^*th*^ column of **X** is distributed as 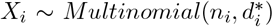, where 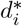 is the *i*^*th*^ column of **D**^*^ [9]. The matrix **D**^*^ satisfies 𝔼[**D**^*^] = **AW**, where **A** ∈ ℝ^*pxK*^ is a non-negative topic matrix, and **W** ∈ ℝ^*Kxn*^ is a non-negative topic weight matrix. The number of topics, *K*, is the dimension of the latent space. Each of the *K* columns of **A** is a probability mass function describing the relative likelihood of each word in the vocabulary with respect to that topic. Each of the *n* columns of **W** describes the mixture of topics within a particular document. By definition, 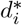 is a convex combination of the columns of **A** weighted and also a probability mass function. The observed corpus matrix **D** is the normalized version of **X** such that each of the columns of **D** sum to 1.

In the context of trajectory inference, we treat the genes as the vocabulary, the cells as the documents, and the expression levels in the corpus matrix as the word counts. The topic matrix, then, is the matrix describing the relationship between the genes and the topics, which we posit correspond to the latent milestones.

Several methods have been proposed to recover the topic matrix from the corpus matrix, including latent Dirichlet allocation [17], non-negative matrix factorization [18], and expectation–maximization algorithm-based methods [19]. Topic-SCORE [9] is a state-of-the-art and computationally efficient method that uses singular value decomposition and simplex geometry to estimate the low-rank **A** from the observed **X**. There are five main steps to Topic-SCORE:

1. **Pre-SVD normalization**. Let **D**^′^ = **M**^—1/2^**D**, where **M** is a diagonal matrix such that **M**_*jj*_ is proportional to the *j*^*th*^ vocabulary word’s frequency in the corpus matrix.
2. **Singular value decomposition**. Calculate Ξ = [*ξ*_1_…*ξ*_*K*_], the *K* left singular vectors of **D**^′^. The rows of Ξ are contained in a simplicial cone with *K* supporting rays. Anchor words (words with nonzero probability on only one topic) lie on the corresponding supporting ray.
3. **Post-SVD normalization**. Perform SCORE normalization [20] on Ξ: normalize each row of Ξ by its first component, yielding **R** ∈ ℝ^*pxK*−1^ and projecting the *K*-dimensional simplicial cone onto a (*K* − 1)-dimensional simplex. Row *j* of **R** is the embedding of word *j* into ℝ^*K*−1^.
4. **Vertex hunting with SVS**. Use Sketched Vertex Search (SVS) [21] to find the vertices of the (*K* − 1)-dimensional simplex. Let 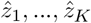 be the vertices found by SVS: these represent the locations of each of the *K* topics in the simplex space. Points within the simplex represent a mixture of topics, and any points outside the simplex (due to noise in the estimation) can be normalized to their nearest point within the bounds of the simplex.
5. **Topic matrix estimation** Let 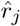 be the *j*^*th*^ row of **R**, which corresponds to the location of word *j* in the (*K* −1)-dimensional simplex. For each vocabulary word, let 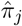 be the coefficients of the convex combination of 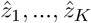 that yields 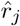. Write 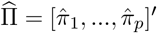. The estimate of **A** is 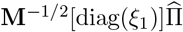.

The topic matrix describes the relationship between the words (genes) and topics (milestones). However, words that are high in frequency across all topics are not helpful for topic interpretation or detection. Therefore, the topic matrix is next converted into a loadings matrix by dividing each row by its *ℓ*^1^-norm. Words corresponding to spiky rows in the loadings matrix—those with one large loading and much smaller values in the remaining components—are more strongly indicative of the topic structure; similarly, genes corresponding to spiky rows in the loadings matrix are more strongly indicative of trajectory- or milestone-relevance. To quantify such trajectory-relevance for each gene, we define a spikiness metric by taking the squared *ℓ*^2^-norm of each row of the loadings matrix. A justification of this metric follows in the next subsection. The spikiness measures are collected into the trajectory relevance vector (TRV).

Crucially, topic modeling requires specifying the number of topics *K*, which cannot be reliably learned from the data. In the trajectory inference, this is equivalent to needing to specify the number of milestones, which is usually unknown. Therefore, to avoid having to select a single number of milestones, we ensemble the TRVs from a range of candidate milestone models using a spectral ensembling procedure. This spectral ensembling method is adapted from the spectral meta-learner of Parisi et al. [10] and the eigenscore method of Ma et al [11]. Let 𝒦 = {*K*_1_, …*K*_|𝒦|_} be the candidate values of *K*, the number of milestones and the number of topics for the topic model. Let *L*_*i*_ be the ℝ^*px*1^ candidate TRV resulting from running the topic model with *K* = *K*_*i*_. We compute a |𝒦| × |𝒦| similarity matrix **G** where **G**_*i,j*_ = *cov*(*L*_*i*_, *L*_*j*_). Let *u* ∈ ℝ^| 𝒦|*x*1^ be the leading eigenvector of **G**. The output of SEEK-VFI is the vector

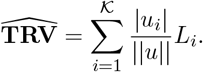

This vector contains the ensembled trajectory relevance score of each gene, which can be used to rank the genes for feature selection and downstream analysis.

### Spikiness metric

Here we will provide a formal argument justifying the proposed spikiness metric. Let the **B** ∈ ℝ^*pxK*^ be the row-normalized gene × milestone matrix, or loadings matrix. By construction, any row **b**_*i*_ = [*b*_*i*,1_, …, *b*_*i,K*_] is such that *b*_*i,j*_ ≥ 0 and ||**b**_*i*_||_1_ = 1, or 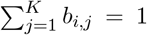. The proposed spikiness measure is 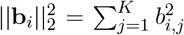. We can set up a Lagrangian with 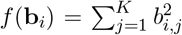 and 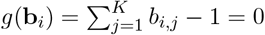:

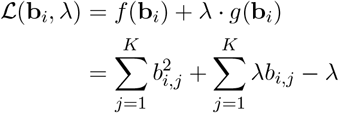

Next, we calculate the gradient:

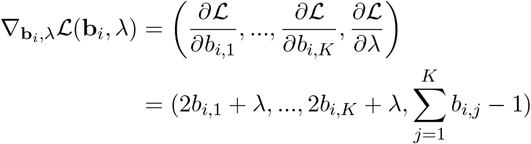

Therefore, we have:

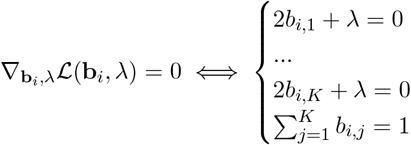

These equations are satisfied (and therefore maxima/minima are achieved) when either of the following is true:

1. λ = 0, *b*_*i,j*_ = 1 for some 1 ≤ *j* ≤ *K*, and *b*_*i,l*_ = 0 for all *l* ≠ *j*
2. 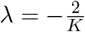 and 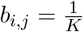 for all 1 ≤ *j* ≤ *K*

Under case 1, which corresponds to the highest possible spikiness setting where a gene is only expressed at one milestone, *f* evaluates to 1. Under case 2, which corresponds to the lowest spikiness setting where a gene is uniformly expressed across milestones, *f* evaluates to 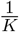. Thus, as desired, our spikiness metric is maximized in the highest-spikiness setting and minimized in the lowest-spikiness setting.

Supplemental Fig. S1 provides an intuitive visual explanation of the metric. These plots show the relationship between the spikiness of a vector **v**_*i*_, with ||**v**_*i*_||_1_ = 1, and its squared *ℓ*^2^-norm 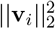 in the *K* = 2 (left) and *K* = 3 (right) cases.

### Simulation details

We tested the performance of SEEK-VFI, along with the three comparator methods from Seurat, on simulated single cell data following six of the basic trajectories described by Saelens et al. [2] and shown in Fig. 2a. The nodes of each trajectory graph represent milestones of cellular development, with individual cells lying on the edges representing transitions between the milestones. We consider two types of genes: trajectory-relevant genes have average expression levels related to the cell’s position in the trajectory (e.g., gene B in Fig 1c), and uninformative genes have constant average expression levels across all locations in the trajectory (e.g., gene A in Fig 1c). Each dataset simulates the expression of 10,000 genes total, with 1,000 trajectory-relevant genes and 9,000 uninformative genes. Each setting comprises 50 simulated datasets.

Let *G* = (*V, E*) denote the trajectory graph, where *v*_1_, …, *v*_*K*_ are the milestones of the graph and edge *e*_*i,j*_ connects *v*_*i*_ and *v*_*j*_. For a gene *g* and milestone *i* ∈ {1, …, *K*}, *p*_*g*_(*i*) is the gene expression probability for a cell at that milestone.

For trajectory-relevant genes, the expression probabilities for a gene *g* are simulated as follows:

1. Let **C** be the distance matrix corresponding to the underlying trajectory: *C*_*ij*_ is the length of the shortest path between milestone *i* and milestone *j*.
2. Define 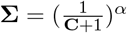.
3. Draw **g**_0_ ∼ *MV N*(0, **∑**).
4. Transform the multivariate normal random vector to a *Uniform*[0, 1] vector via **g**_1_ = Φ(**g**_0_), where Φ is the normal CDF.
5. Linearly rescale **g**_1_ to **g** ∼ *Uniform*[0.2, 0.8]; this ensures that the resulting cell counts are reasonable.
6. The expression probability for gene *g* with respect to milestone *i* is *p*_*g*_(*i*) = **g**_*i*_, the *i*^*th*^ entry of **g**. The vector **g** is analogous to the *g*^*th*^ row of the topic matrix in the standatd topic modeling framework.

In other words, for trajectory-relevant genes, the expression probability is indexed by the milestone, and the probabilities are correlated with respect to the distance between nodes in the underlying trajectory.

For uninformative genes, the expression probability is constant across nodes: *p*_*g*_(*i*) ∼ *Uniform*[0.2, 0.8] and *p*_*g*_(*i*) = *p*_*g*_(*j*) for all milestones *i* and *j*.

Suppose a cell *c* is *d*% along the transition from milestone *v*_*i*_ to milestone *v*_*j*_: *c* lies on *e*_*i,j*_, its scaled distance from *v*_*i*_ is *d*, and its scaled distance from *v*_*j*_ is 1 *— d*. For a gene *g*, the gene expression probability for that cell is *p*_*c,g*_ = *d* · *p*_*g*_(*i*) + (1 *— d*) · *p*_*g*_(*j*). Draw a value *r* ∼ *N*(2, 0.5), and let *r*_*g*_ = *max*(*r*, 0.1) to yield a gene-specific baseline expression level. Then the expression of gene *g* for cell *c* is drawn from *NegBinomial*(*r*_*g*_, *p*_*c,g*_).

For each setting (trajectory type, number of cells, and *α*), we generate 50 datasets representing simulated gene expression from the given number of cells.

### Real-data analysis details

The real datasets analyzed in this paper are:

- **Fibroblast:** bifurcation trajectory representing the reprogramming of mouse embryonic fibroblasts into myocytes and neurons [22]
- **Hematopoiesis:** complex tree structure of hematopoietic differentiation [23]
- **Kidney:** linear trajectory of kidney duct cells through a transitional cell type [24]
- **Oligodendrocytes:** linear progression of oligodendrocyte precursor cells to mature oligodendrocytes [25]
- **Thymus:** bifurcation trajectory of t-cells in the thymus [26]

For the analysis in Fig. 1b, the ground-truth distance matrices were derived using the cell labels and milestone networks provided in the data [27]. The ground-truth distance between two cells is defined as the length of shortest path between the two cell states in the milestone network.

We used the default settings for UMAP and t-SNE to generate the low-dimensional visualizations in Fig. 1c.

For analysis with Monocle3, Slingshot, and Totem, we provided as input data the counts matrix restricted to the top 500 genes from each method and the UMAP embedding derived from the top 500 genes as the reduced dimensions object. We ran Slingshot via *dynverse* [2]. For all methods, we provided a start cell by selecting the median point within the appropriate cell state with respect to the low-dimensional embedding.

To evaluate the performance of the downstream analysis methods, we consider a) the estimated pseudotime for each cell and b) the estimated milestone network. To evaluate the milestone network, we assign each cell to a cluster based on its nearest milestone and compare it to the ground-truth cell state using the normalized mutual information (Fig. 2d, x-axis). To evaluate the pseudotime, we calculate its correlation to the pseudotime derived from the ground-truth milestone network (Fig. 2d, y-axis). The ground-truth milestone network pseudotime is calculated as follows:

Let **G** = {**V, E**} be the directed graph corresponding to the ground truth milestone network. Let *v*_0_ ∈ **V** be the root of the graph and let **V**^*^ ⊂ **V** be the set of leaf nodes: *indegree*(*v*_0_) = 0 and *outdegree*(*v*^*^) = 0 for *v*^*^ ∈ **V**^*^. For each *v*^*^ ∈ **V**^*^, let **G**[*v*^*^] = {**V**[*v*^*^], **E**[*v*^*^]} be the subgraph of **G** corresponding to the shortest path between *v*_0_ and *v*^*^. For *v* ∈ *V* [*v*^*^], define 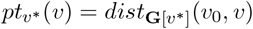, the distance between the root node and the node of interest with respect to the subgraph. If *v* is not in ***G***[*v*^*^], we set the value as null and remove it prior to the final pseudotime calculation. The ground-truth milestone network pseudotime for a node *v* ∈ **V** is 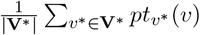, or the average pseudotime across all subgraphs.

### Hyperparameter selection and computational cost

A key hyperparameter for SEEK-VFI is the choice of candidate milestone counts *K* to ensemble. Underspecified models (chosen *K* is smaller than the truth) capture a subset of the signal, while overspecified models (chosen *K* is greater than the truth) result in signal + noise [9]. The ensembling method is highly robust to the inclusion of noise [11], and in simulations experiences minimal degradation in quality when additional overspecified models are included. However, there is a tradeoff regarding computational time: while Topic-SCORE is much faster than other topic modeling methods [9], its most computationally intensive step (Sketched Vertex Search for vertex hunting) requires iterating through 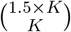 combinations of candidate vertices, which accumulates exponentially as *K* increases (Supplemental Fig. S2a). While the simulations and real-data results in this paper were run using SEEK-12, since the computational time is very reasonable given the data size, the results in Supplemental Fig. and show that the results for SEEK-6 are quite similar to those in the main figure, and that reduced upper ensembling limits can be chosen if faster computational time is desired.

## Data and code availability

The code for running SEEK-VFI is available to download as an R package at github.com/rdanning/seekvfi. The real datasets are available at zenodo.org/records/1443566 [27].

## Acknowledgements

R.D.’s work was supported by NIH Grant T32-GM135117 while at the Harvard TH Chan School of Public Health. Z.T.K.’s work is supported by the Sloan Research Grant FG-2023-19970. X.L.’s work is supported by NIH Grants R35-CA197449, R01-HL163560, and U01-

