## Supplemental Material for "Trajectory-informed gene feature selection in single-cell analysis with SEEK-VFI"

### Supplemental information

#### Methods figures

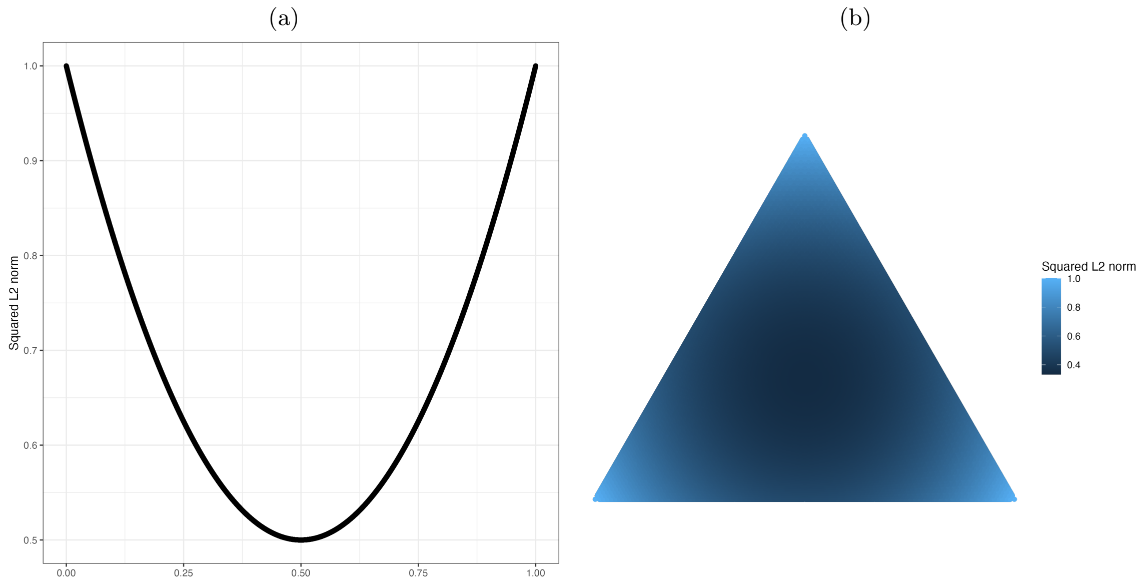


Supplemental Figure S1: (a) Plot comparing the relationship between vector elements and squared $\ell^2$-norm for vectors of 2 elements that sum to 1. The x-axis represents the value of the first coordinate, and thus the other coordinate is equal to 1 minus the first coordinate. The y-axis represents the value of the squared $\ell^2$-norm. Points at the edges of the x range represent spikier vectors, whereas points in the middle represent vectors where the weight is distributed more evenly across the two coordinates. (b) Plot comparing the relationship between vector elements and squared $\ell^2$-norm for vectors of 3 elements that sum to 1. Points at the vertices of the triangle represent vectors with all of their weight in just one element, whereas points towards the middle represent vectors with an even distribution of weights across the three elements.

#### Seurat comparator methods

The dispersion method ("Disp") directly calculates the dispersion of each gene by computing the ratio of the sample variance to the sample mean with respect to the log-transformed expression counts [3]. The scaled dispersion method ("ScDisp") normalizes the dispersion scores with respect to bins of cells with similar average expression [6]. The variance-stabilizing method ("VST") transforms the counts into Z-scores using a variance-stabilizing transformation and then calculates the variance with respect to these (potentially truncated) Z-scores to assess the dispersion of the expression counts while also controlling for the average expression of each gene [7].

#### SEEK-6 vs. SEEK-12

#### Additional results

Supplemental Fig. S4 shows the variation in performance across trajectory types for each method. In this plot, the $x$-axis represents the number of cells, while $\alpha$ is fixed at 0.25. SEEK-VFI is the most robust to trajectory type, with consistent performance once the sample size is at least 1000 cells. The Seurat methods, on the other hand, experience worse performance with the multifurcation and tree settings, even when the number of cells is larger. Supplemental Fig. S5 provides expanded results from Fig. 1b; notably, as the percentage of genes selected increases, the methods become more similar, indicating that they are identifying similar gene lists.

##### Comparison with DELVE

DELVE is a recently-developed method for unsupervised feature selection in molecular data that is designed to capture features most relevant to the underlying cellular trajectory through the identification of modules of features that vary significantly across inferred cell states [12]. Both SEEK-VFI and DELVE model the underlying trajectory structure prior to feature ranking; while SEEK-VFI uses SVD-based methods to model the latent space, DELVE models the cellular trajectory via a k-nearest neighbor affinity graph where the edge weight between cells is a function of similarity across all observed features. In our initial simulations, we tested DELVE alongside SEEK-VFI and the Seurat methods and found its performance comparable to the Seurat methods (Supplemental Fig. S3). Likewise, DELVE took around 2 hours and 45 minutes to run in settings where the other methods took less than 3 minutes. As such, we chose to exclude DELVE from further analyses.

##### Additional low-dimensional visualizations

While the results in Fig. 2c and d are based on the selection of the top 500 genes from each method, and the low-dimensional embedding from UMAP, the following figures show visual results comparing the methods across the top 250, 500, 750, and 1000 genes, as well as across UMAP, t-SNE, and PCA.

(a)

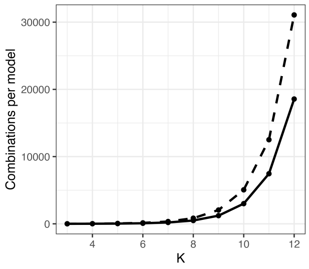


(b)

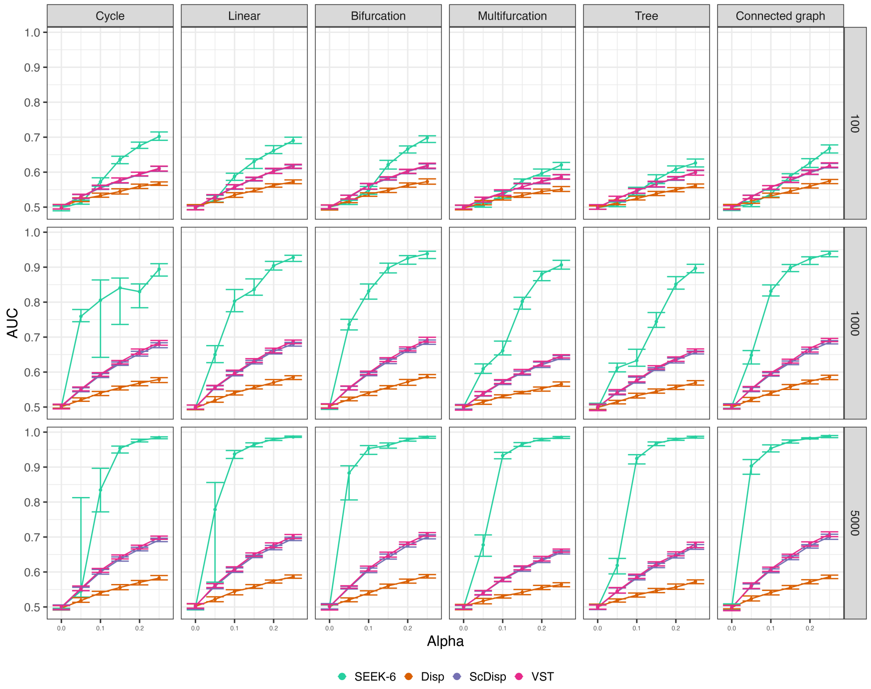


(c)

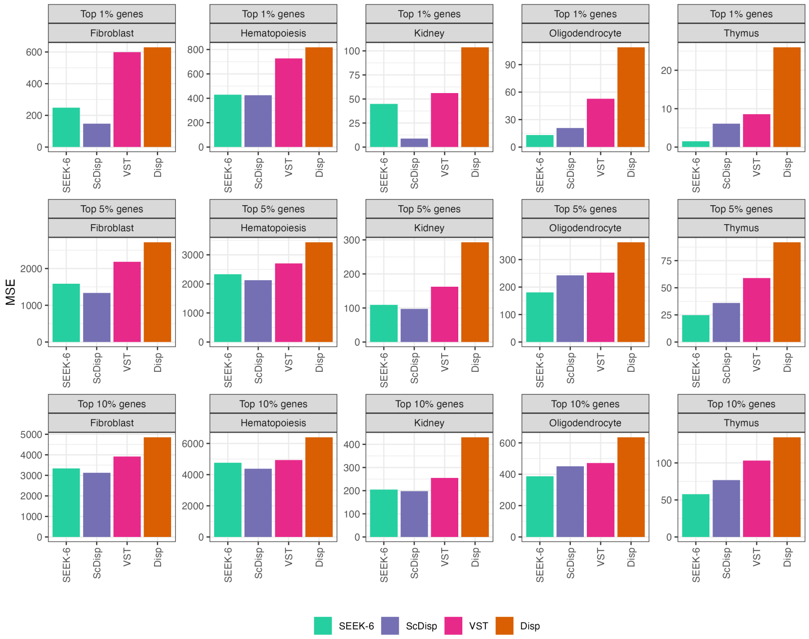


Supplemental Figure S2: (a) Solid line: plot of $K$ versus $\binom{1.5\times K}{K}$ showing the number of vertex combinations considered in the Sketched Vertex Search step of Topic-SCORE for a model with $K$ topics. Dashed line: cumulative sum showing the total number of combinations required when running models $3, ..., K$ to ensemble. (b) Simulated data results analogous to Fig. 1a with SEEK-6 (ensembling $K = \{3, ...6\}$) instead of SEEK-12 (ensembling $K = \{3, ...12\}$). (c) Real-data results analogous to Fig. 1b with SEEK-6 instead of SEEK-12.

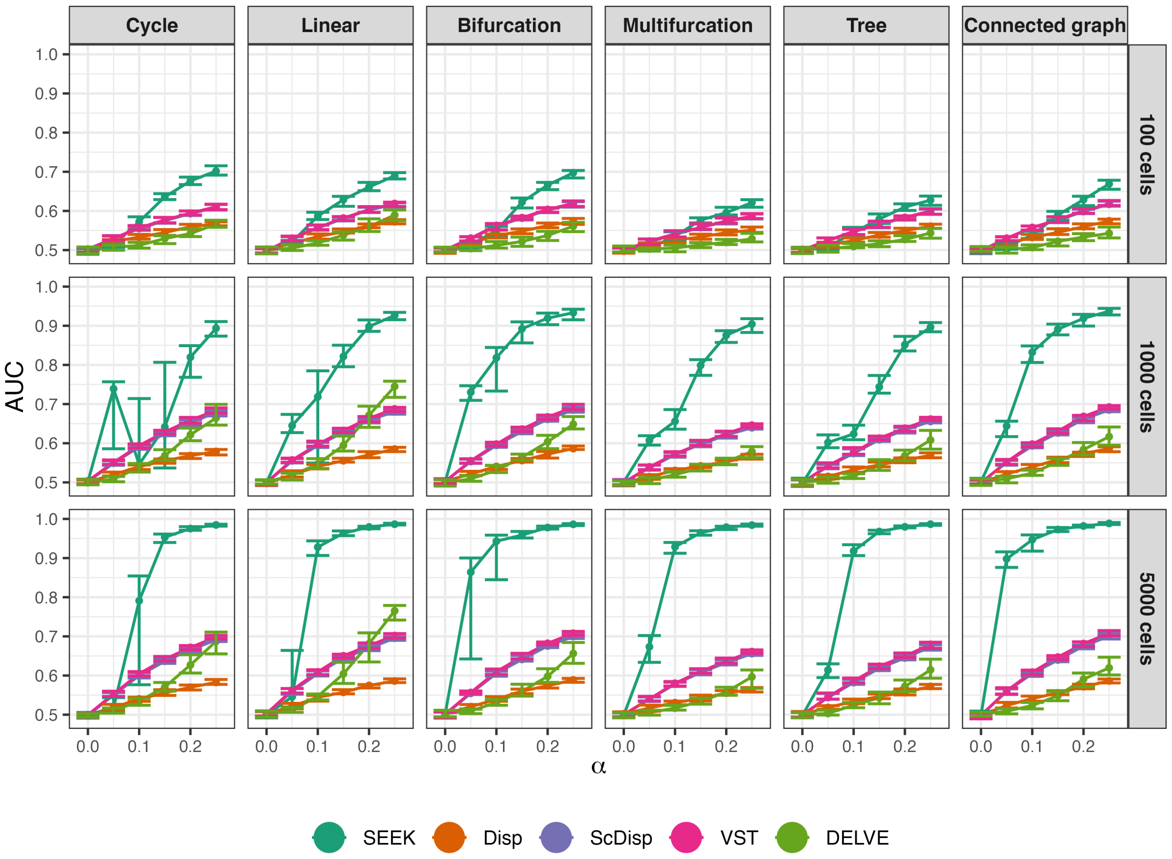


Supplemental Figure S3: Expanded version of Fig. 2a showing results with a varied number of cells. Fewer cells represents a more challenging identification task.

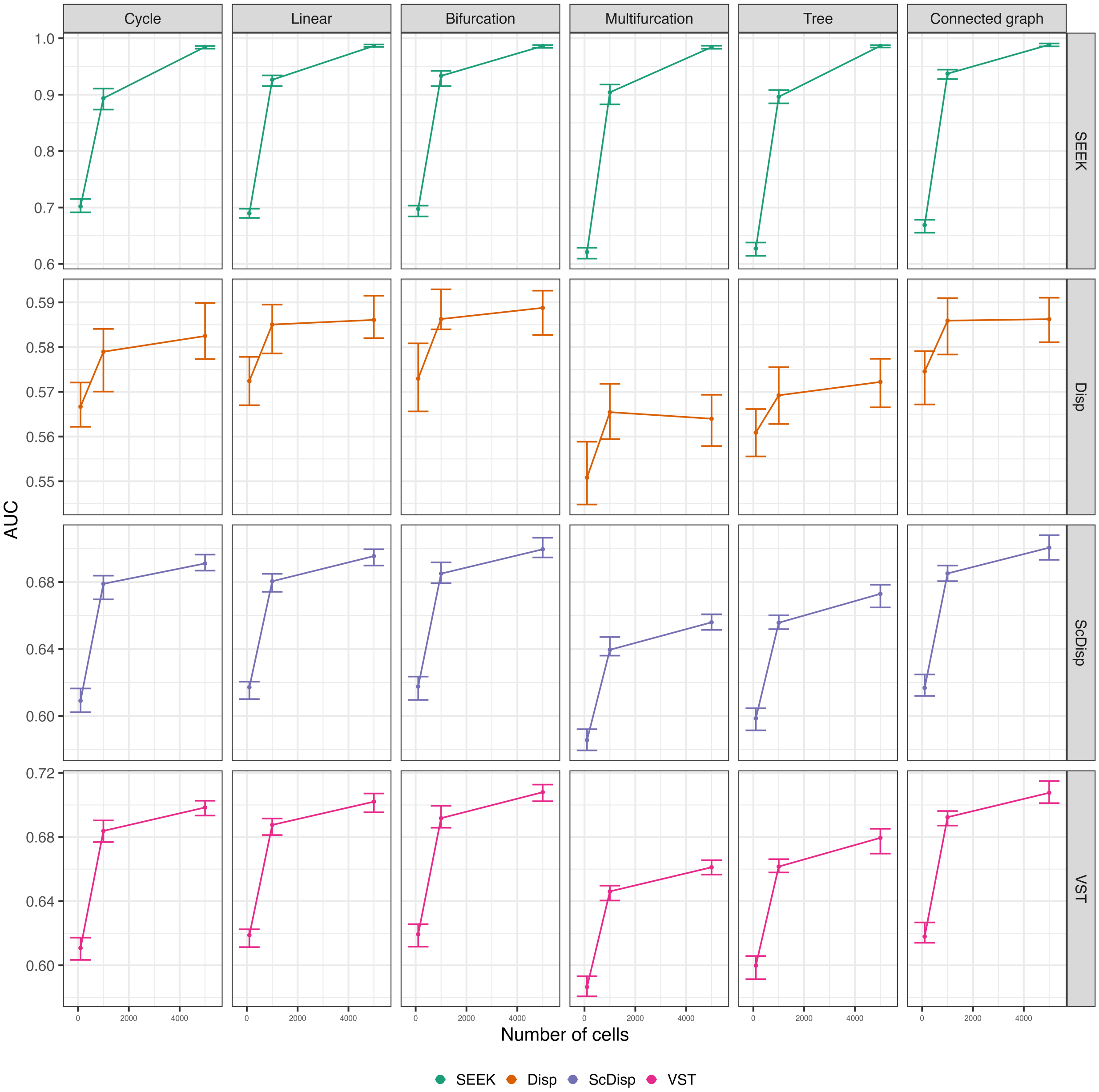


Supplemental Figure S4: Performance of SEEK-VFI and the three Seurat methods on simulated data with 50 simulated datasets per setting. Columns correspond to trajectory types and rows separate the results for each method. The value of $\alpha$ is set at 0.25, and the x-axis varies the number of cells. The y-axis shows the median AUC for differentiating between trajectory-relevant and uninformative genes across the simulations. Error bars represent the $25^{th}$ and $75^{th}$ percentiles.

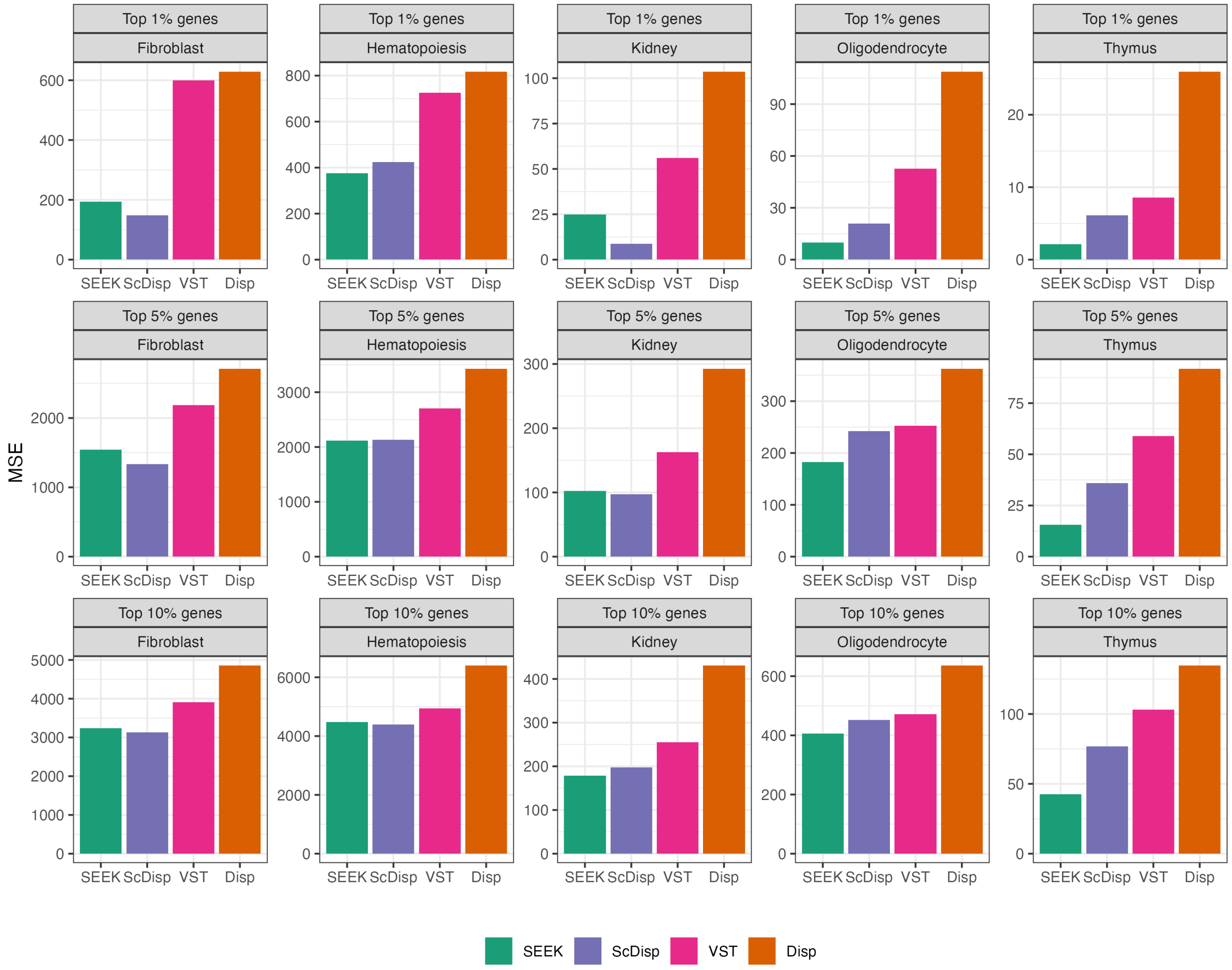


Supplemental Figure S5: Bar plots showing the mean squared error comparing the true trajectory distance matrix to the distance matrices calculated with respect to expression of the top X% of genes as selected by each of the four comparator methods.

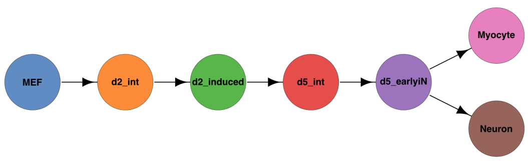


Supplemental Figure S6: Plot of the fibroblast ground-truth trajectory structure.

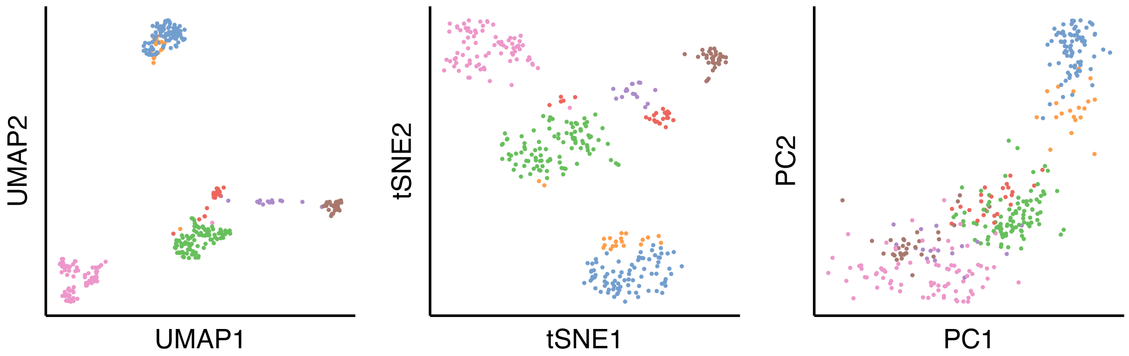


Supplemental Figure S7: Plots of the UMAP, t-SNE, and PCA embeddings of the fibroblast cells using all 3379 genes.

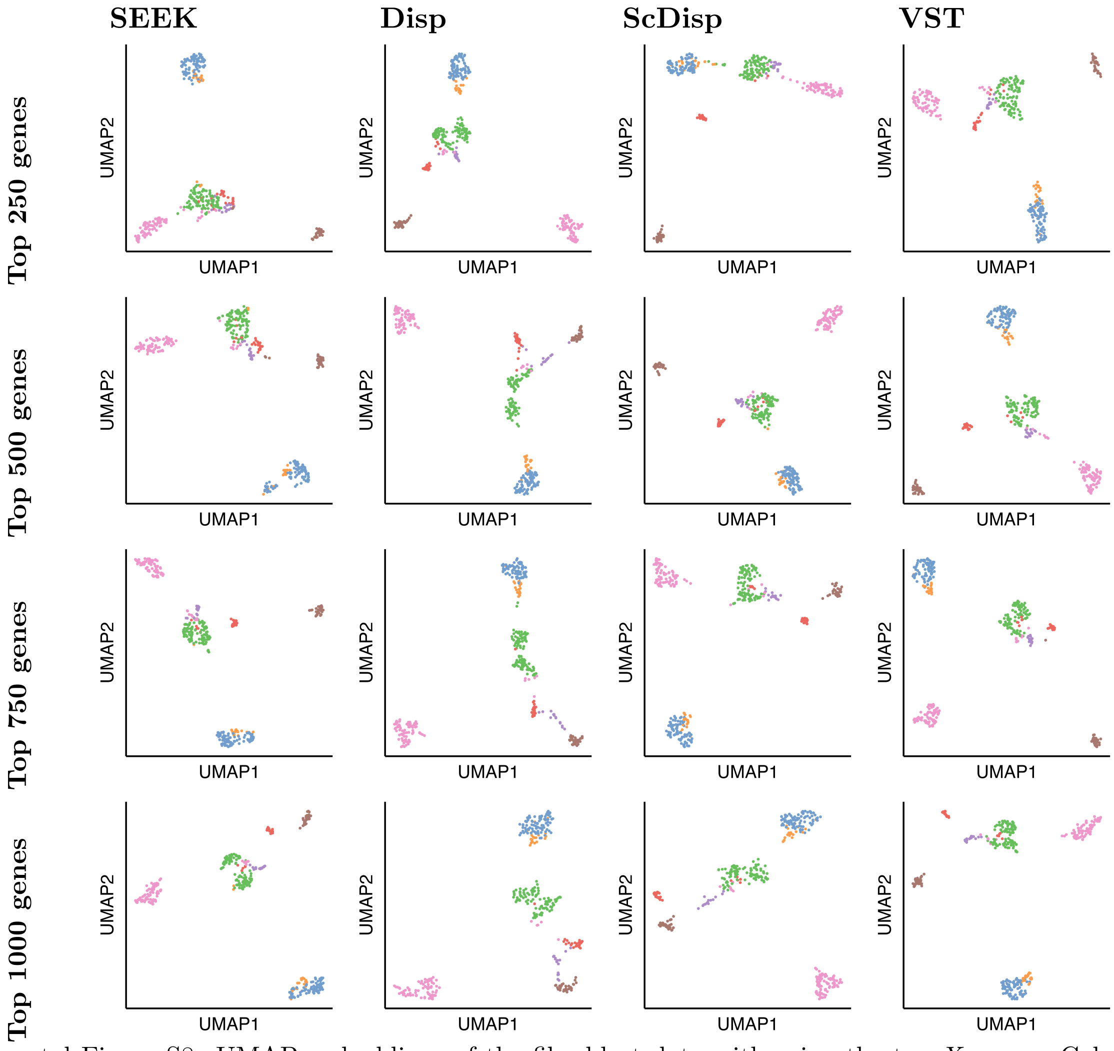


Supplemental Figure S8: UMAP embeddings of the fibroblast data with using the top $X$ genes. Columns correspond to the method used to select the top genes and rows correspond to the value of $X$.

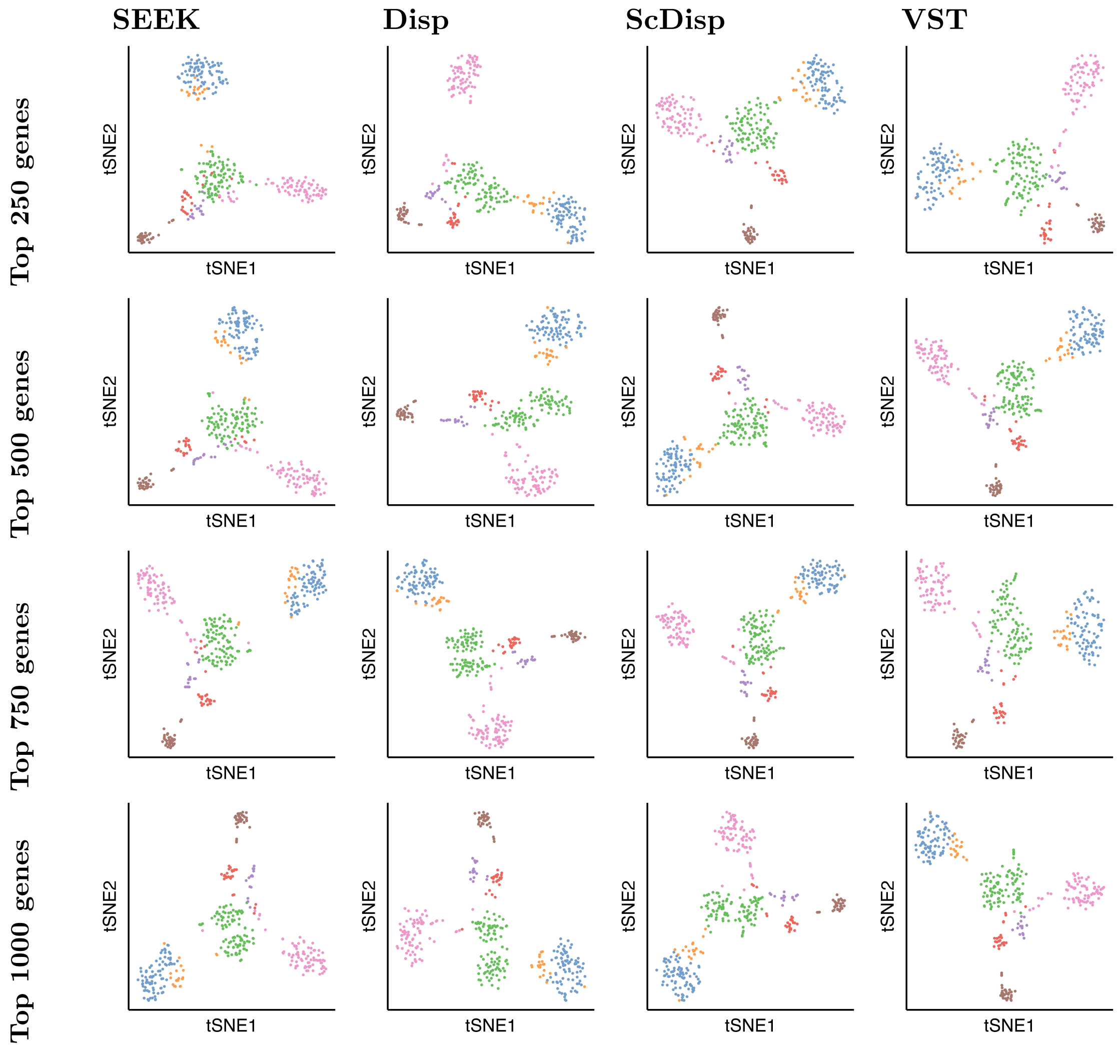

Supplemental Figure S9: t-SNE embeddings of the fibroblast data with using the top $X$ genes. Columns correspond to the method used to select the top genes and rows correspond to the value of $X$.

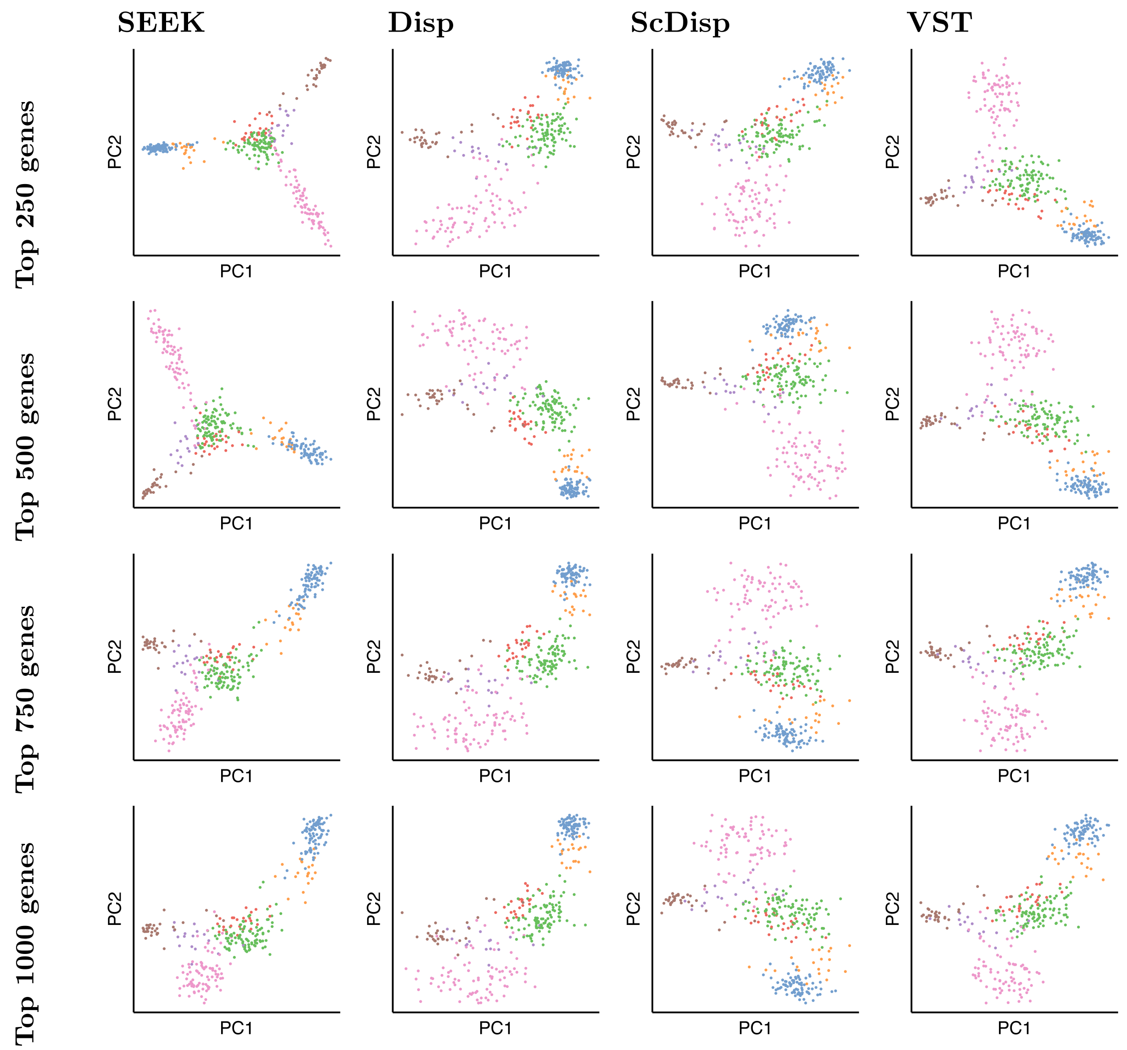

Supplemental Figure S10: PCA embeddings of the fibroblast data with using the top $X$ genes. Columns correspond to the method used to select the top genes and rows correspond to the value of $X$.

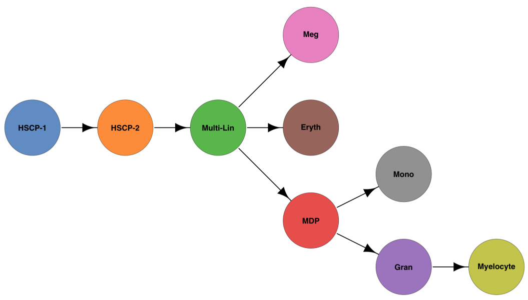


Supplemental Figure S11: Plot of the hematopoiesis ground-truth trajectory structure.

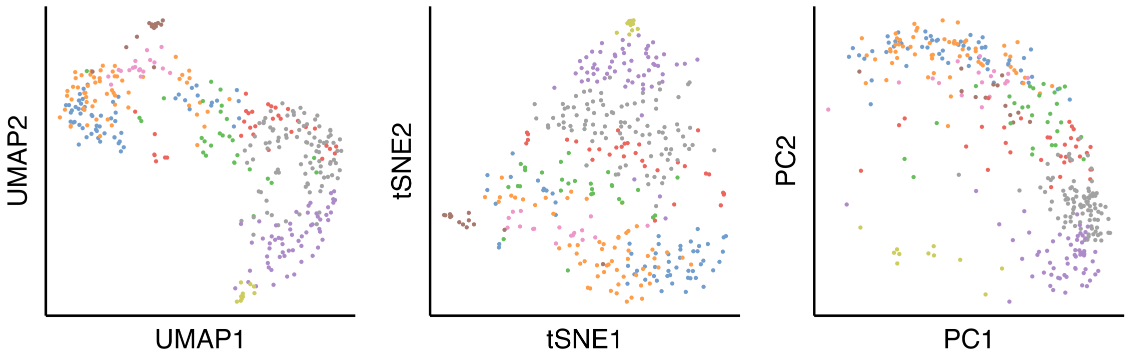


Supplemental Figure S12: Plots of the UMAP, t-SNE, and PCA embeddings of the hematopoiesis cells using all 3594 genes.

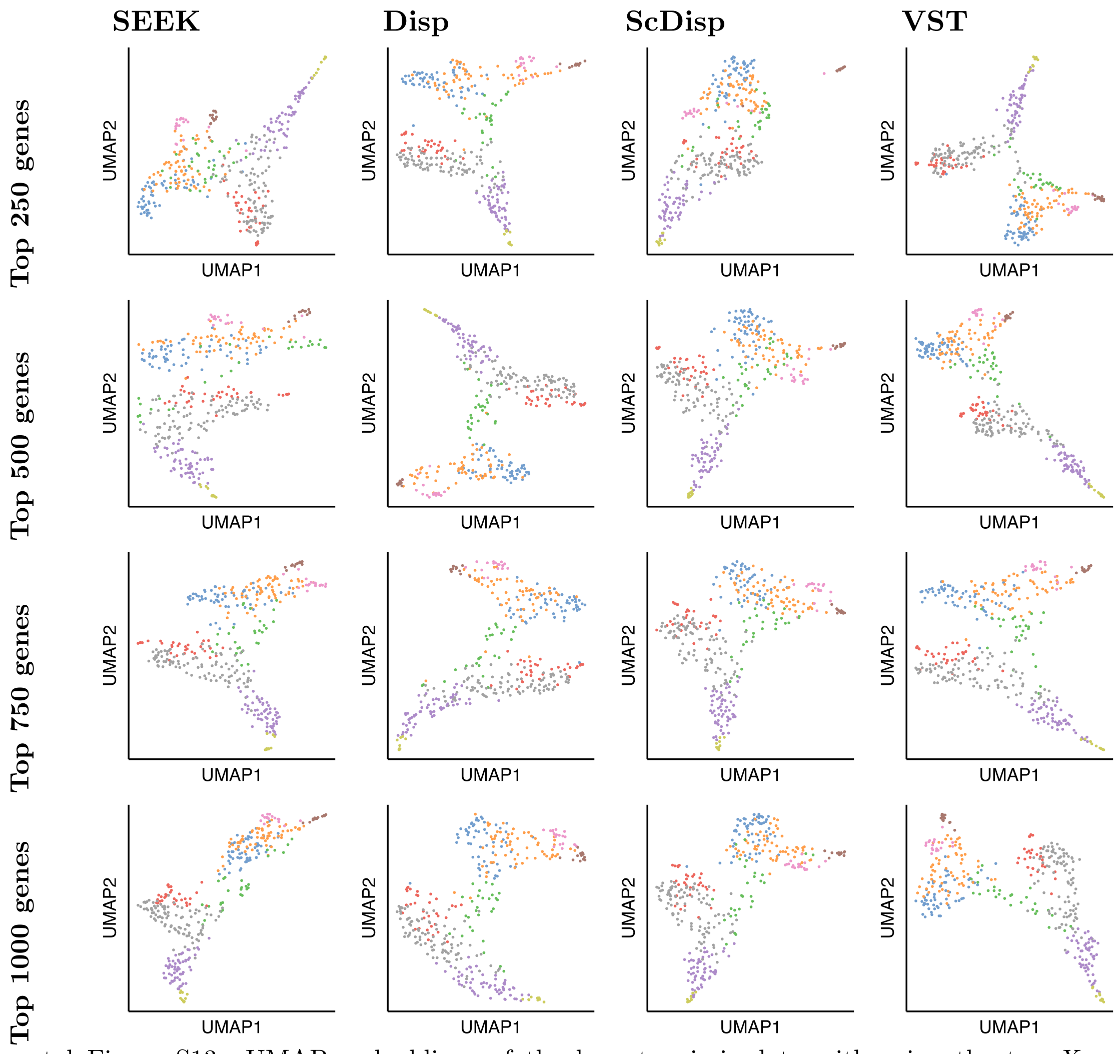


Supplemental Figure S13: UMAP embeddings of the hematopoiesis data with using the top $X$ genes. Columns correspond to the method used to select the top genes and rows correspond to the value of $X$.

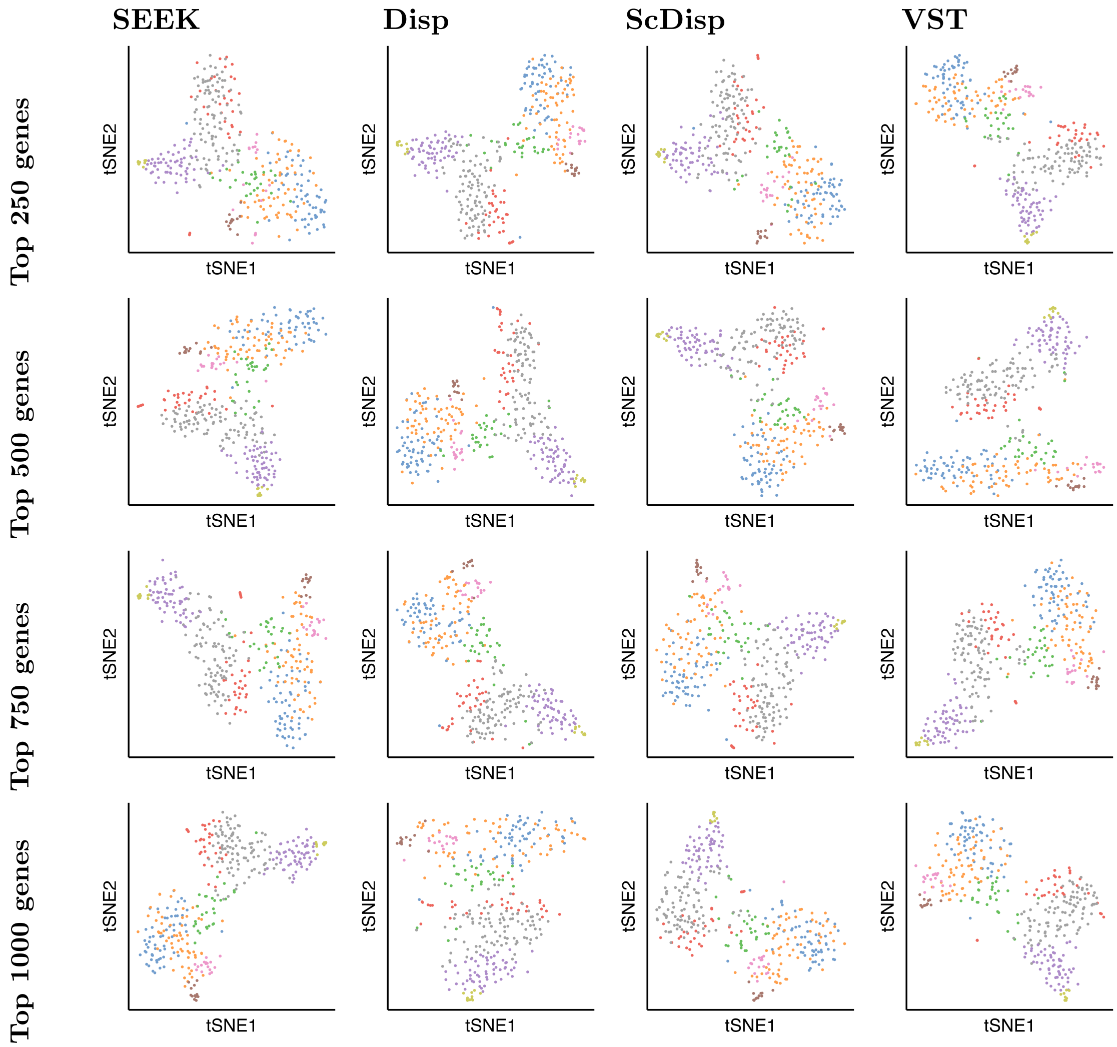


Supplemental Figure S14: t-SNE embeddings of the hematopoiesis data with using the top $X$ genes. Columns correspond to the method used to select the top genes and rows correspond to the value of $X$.

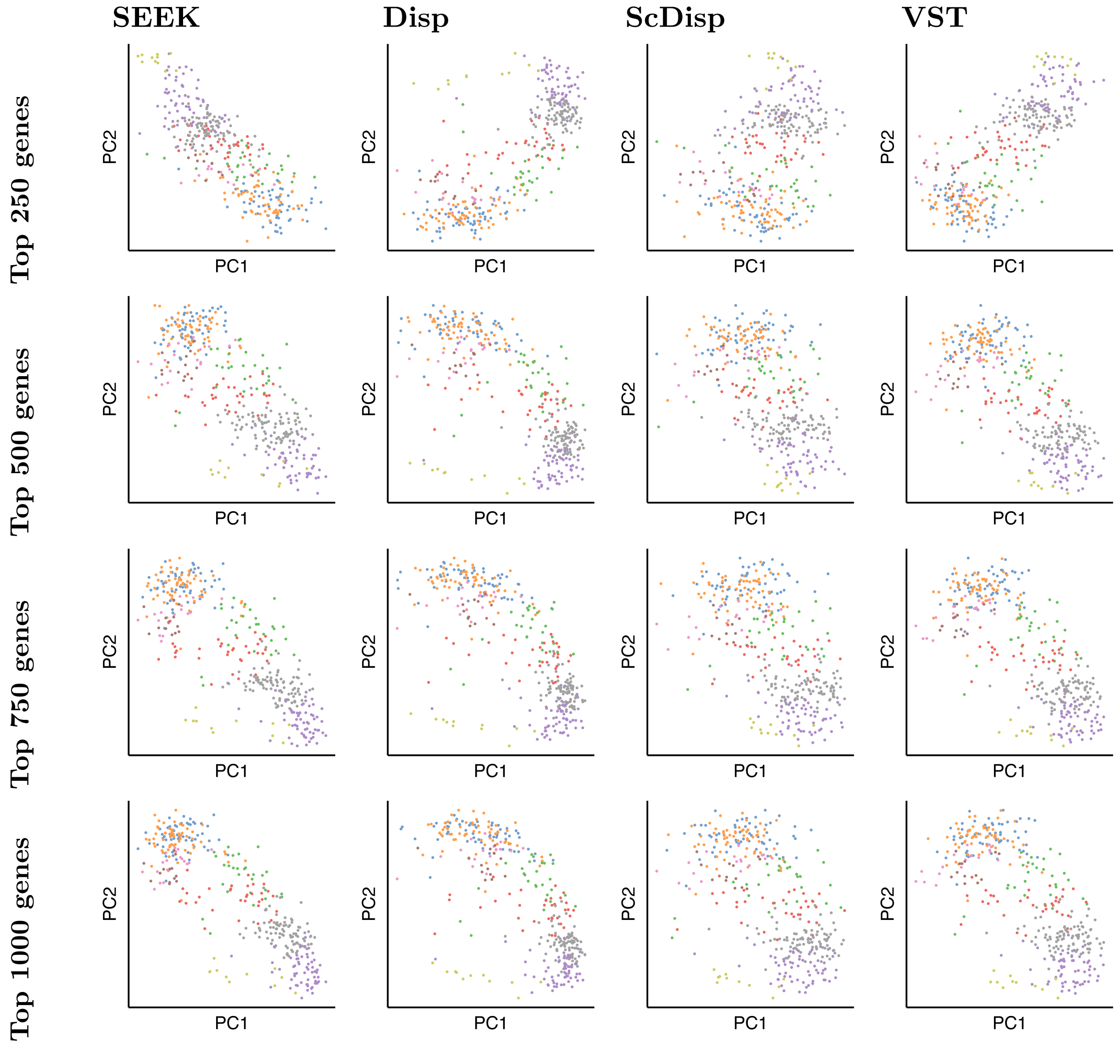


Supplemental Figure S15: PCA embeddings of the hematopoiesis data with using the top $X$ genes. Columns correspond to the method used to select the top genes and rows correspond to the value of $X$.

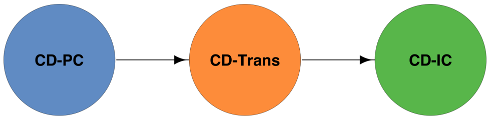


Supplemental Figure S16: Plot of the kidney ground-truth trajectory structure.

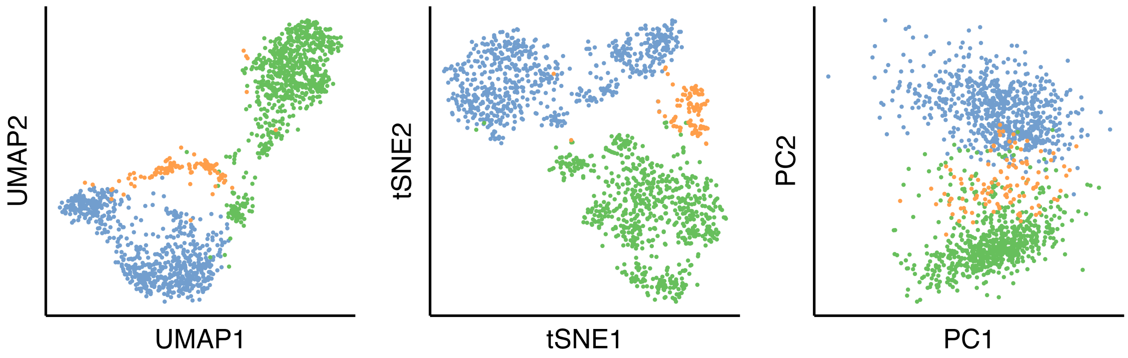


Supplemental Figure S17: Plots of the UMAP, t-SNE, and PCA embeddings of the kidney cells using all 2441 genes.

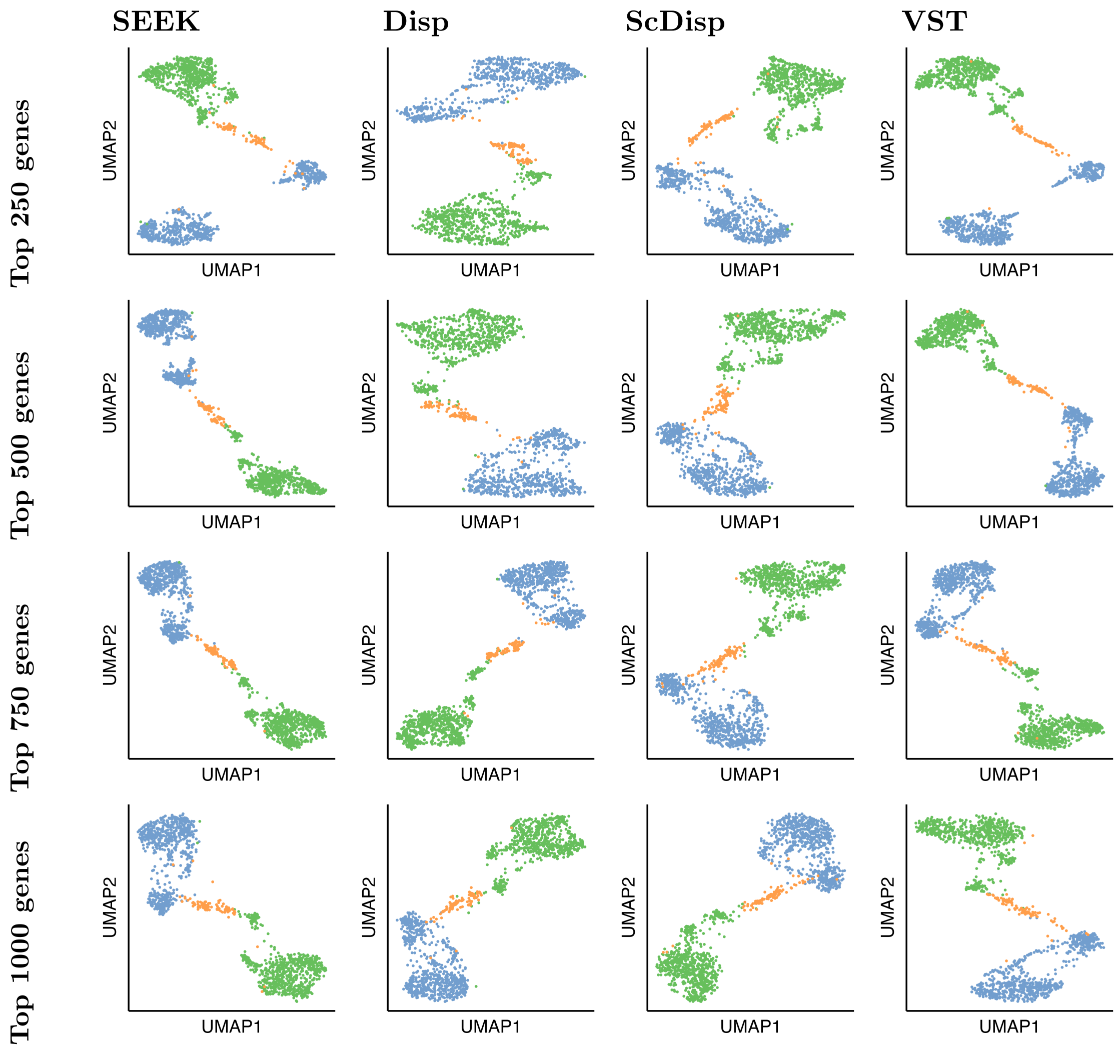


Supplemental Figure S18: UMAP embeddings of the kidney data with using the top $X$ genes. Columns correspond to the method used to select the top genes and rows correspond to the value of $X$.

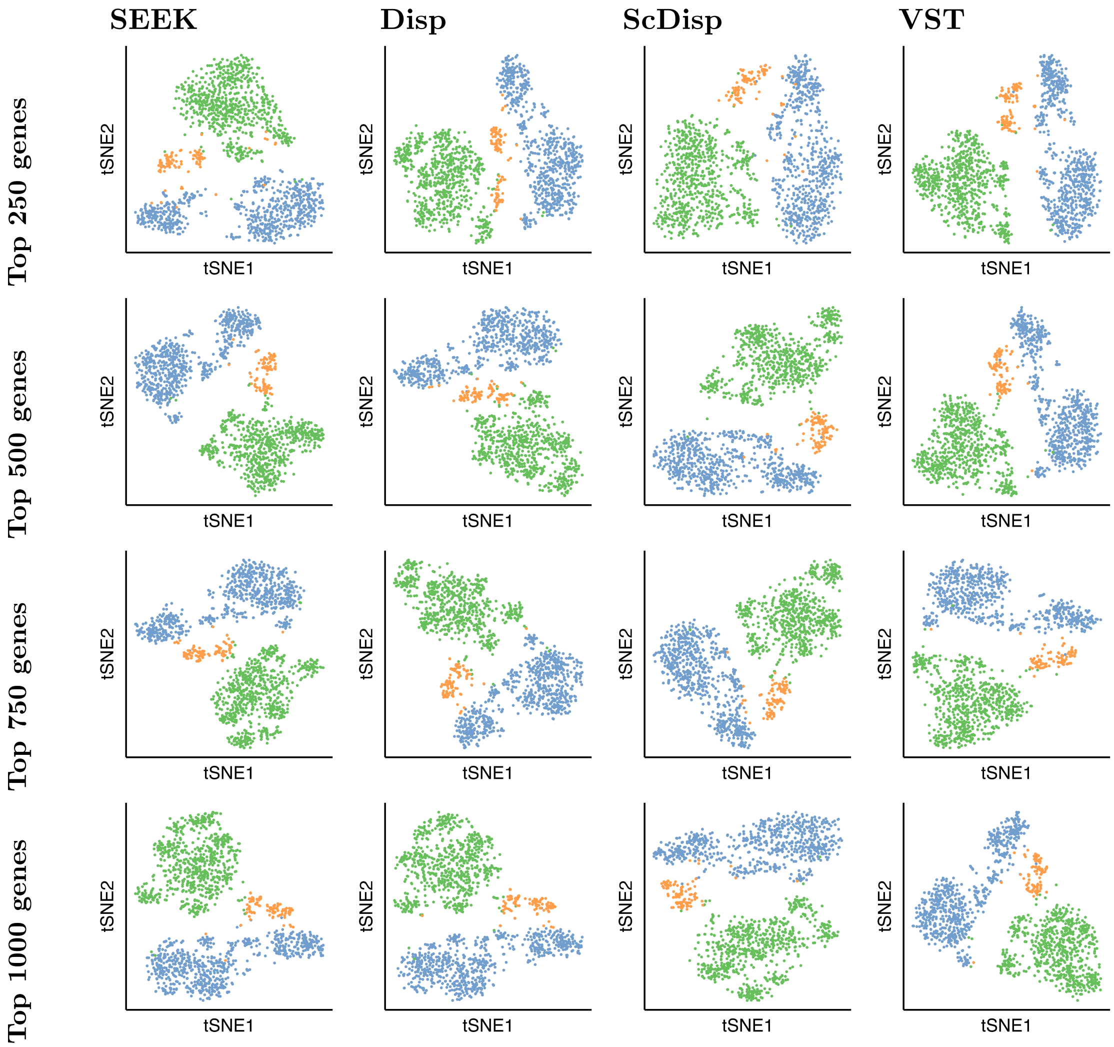

Supplemental Figure S19: t-SNE embeddings of the kidney data with using the top $X$ genes. Columns correspond to the method used to select the top genes and rows correspond to the value of $X$.

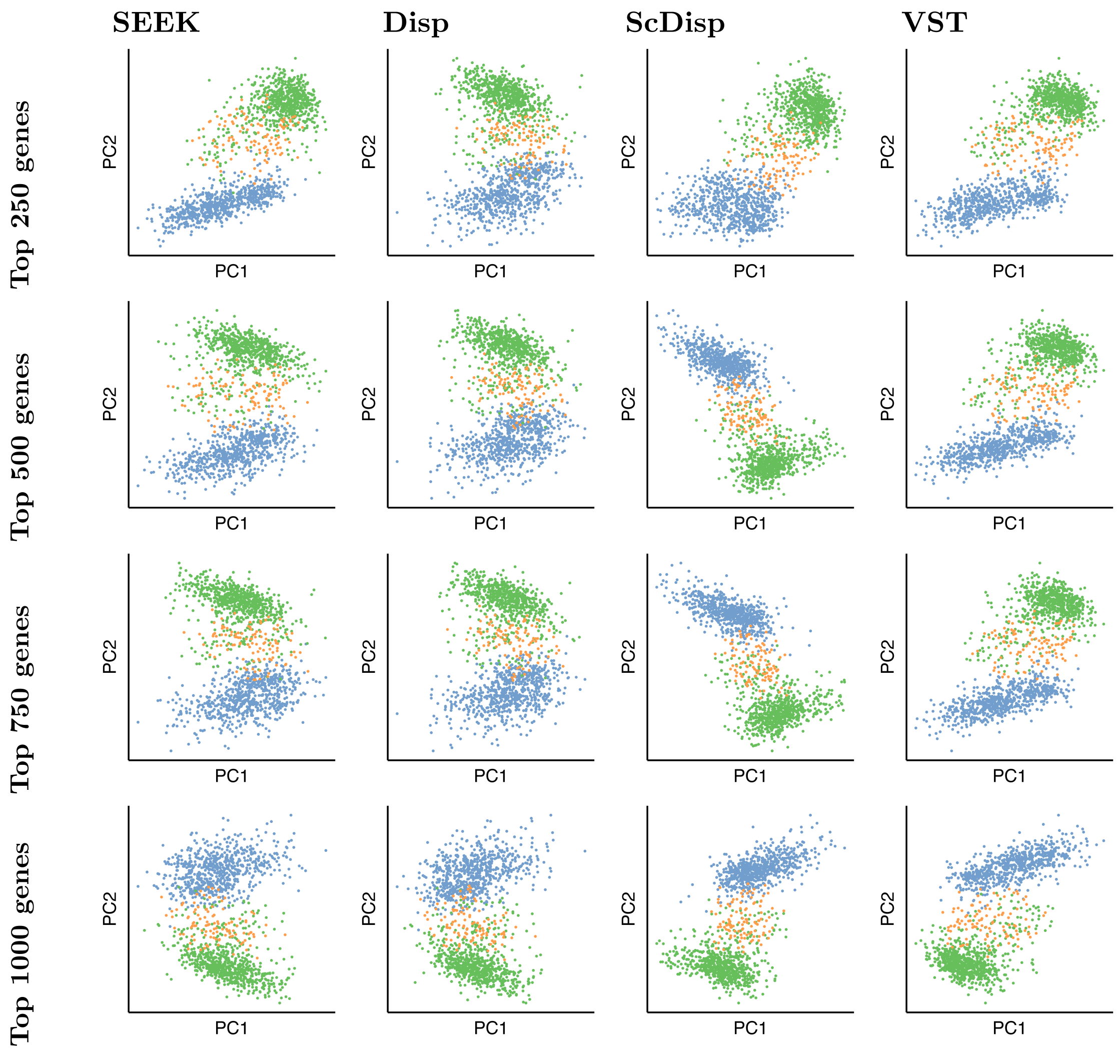


Supplemental Figure S20: PCA embeddings of the kidney data with using the top $X$ genes. Columns correspond to the method used to select the top genes and rows correspond to the value of $X$.

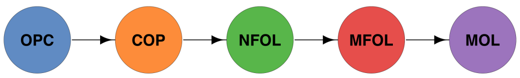


Supplemental Figure S21: Plot of the oligodendrocyte ground-truth trajectory structure.

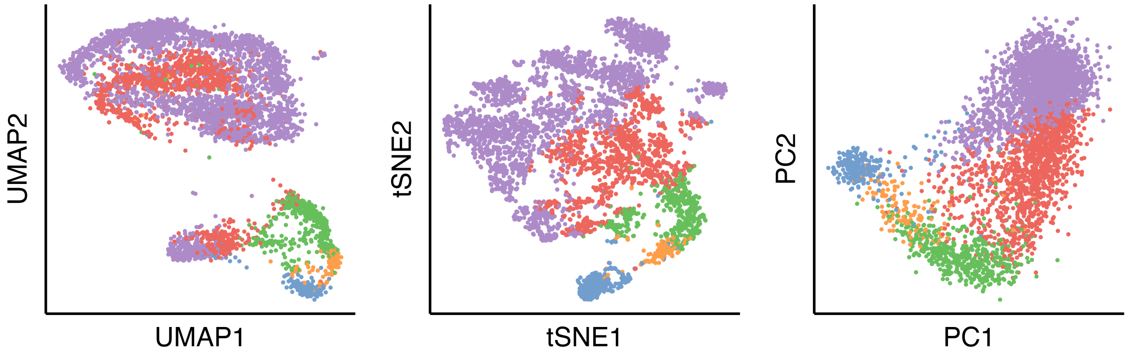


Supplemental Figure S22: Plots of the UMAP, t-SNE, and PCA embeddings of the oligodendrocyte cells using all 3534 genes.

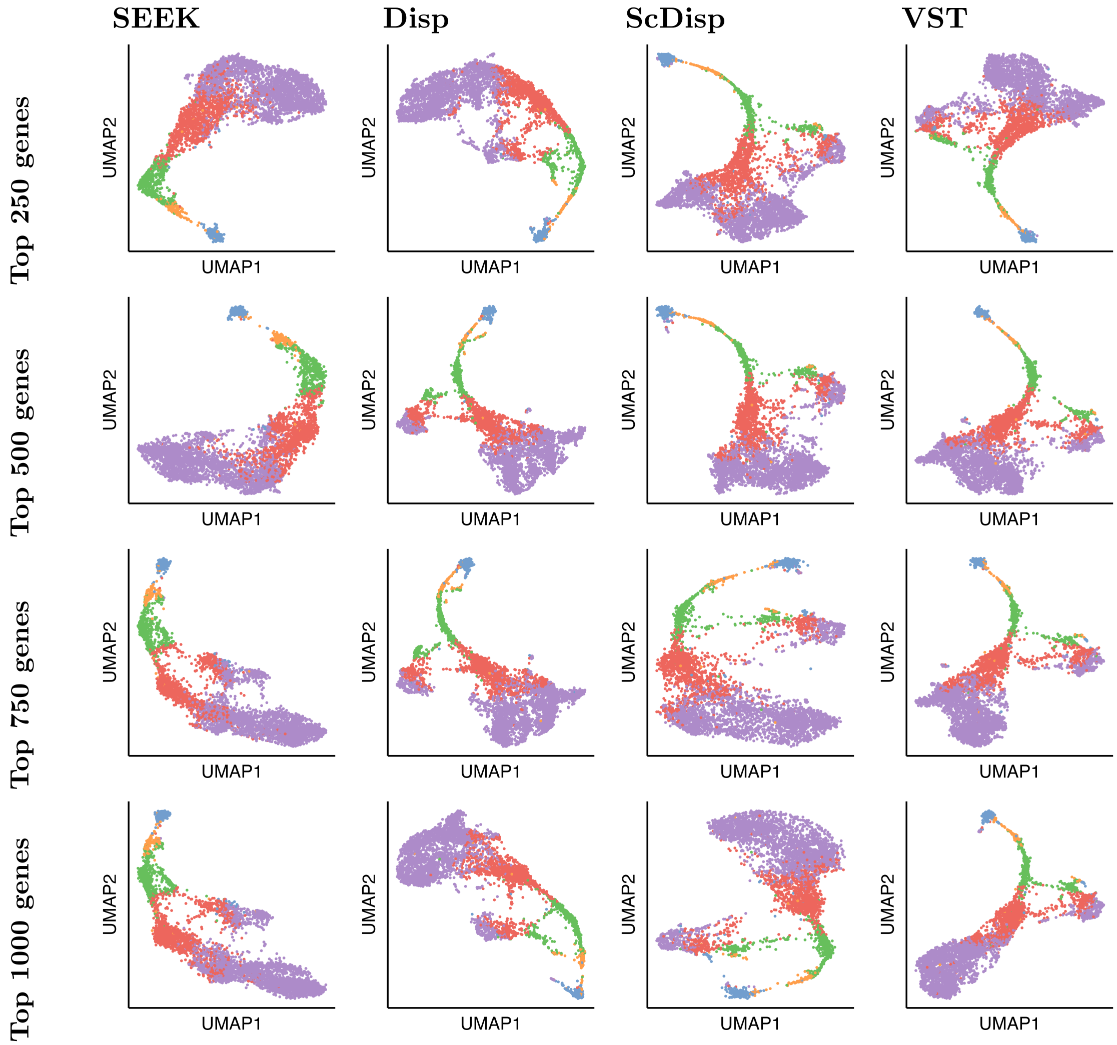


Supplemental Figure S23: UMAP embeddings of the oligodendrocyte data with using the top $X$ genes. Columns correspond to the method used to select the top genes and rows correspond to the value of $X$.

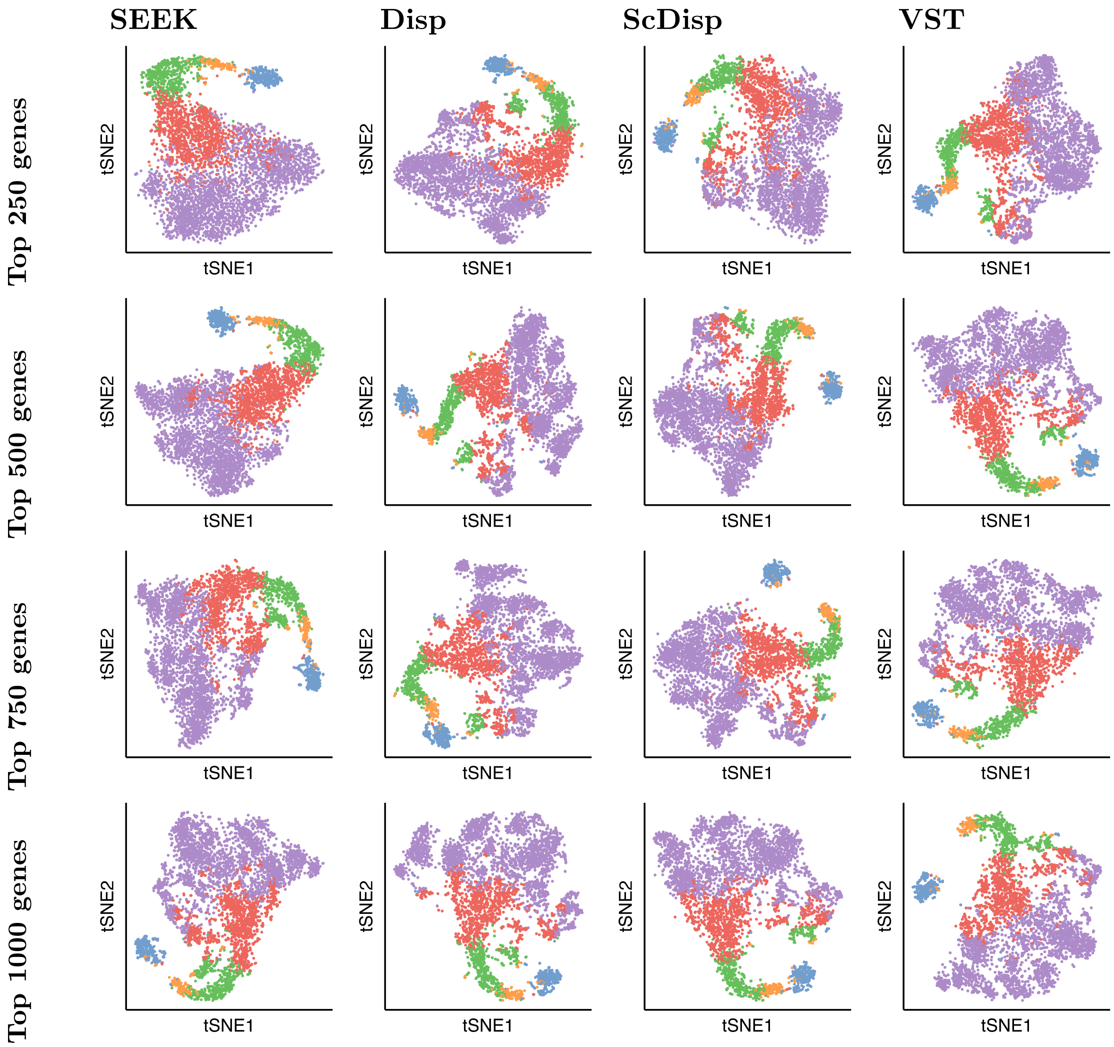


Supplemental Figure S24: t-SNE embeddings of the oligodendrocyte data with using the top $X$ genes. Columns correspond to the method used to select the top genes and rows correspond to the value of $X$.

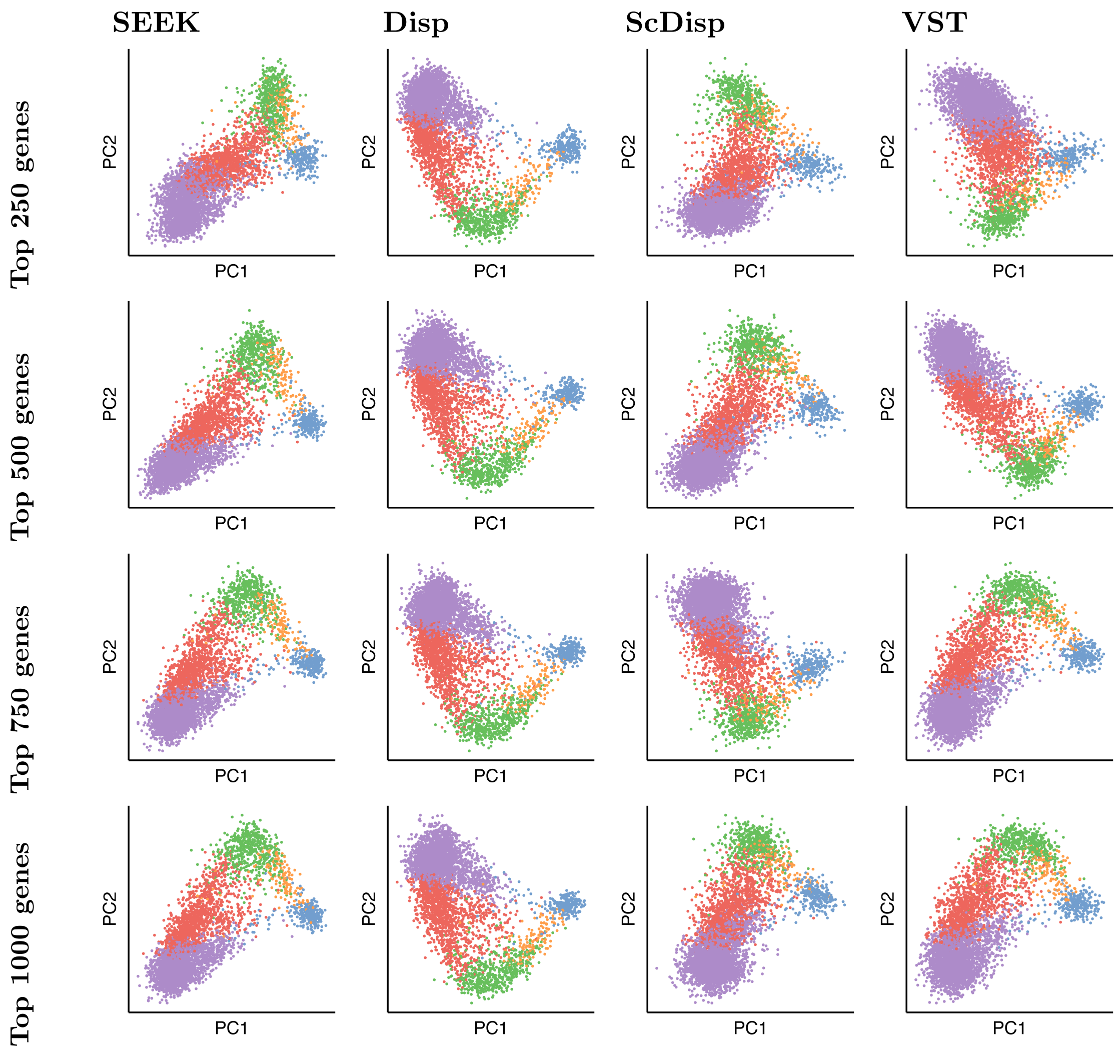

Supplemental Figure S25: PCA embeddings of the oligodendrocyte data with using the top $X$ genes. Columns correspond to the method used to select the top genes and rows correspond to the value of $X$.

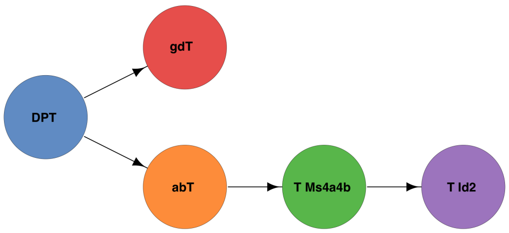

Supplemental Figure S26: Plot of the thymus ground-truth trajectory structure.

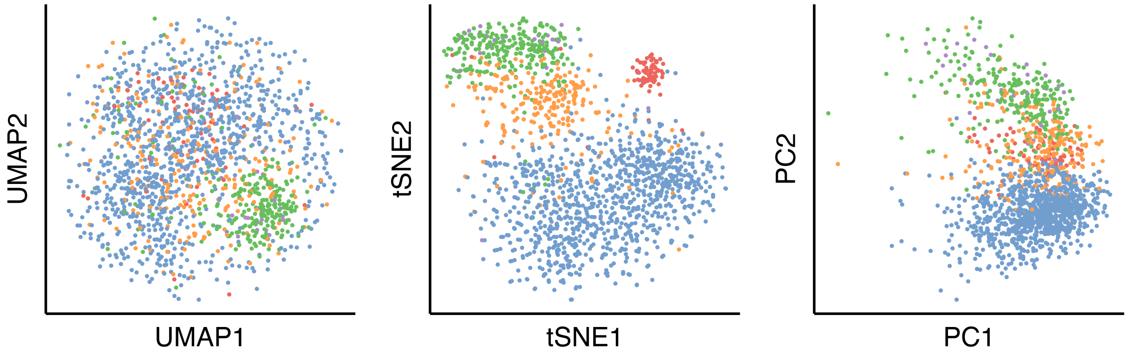


Supplemental Figure S27: Plots of the UMAP, t-SNE, and PCA embeddings of the thymus cells using all 2642 genes.

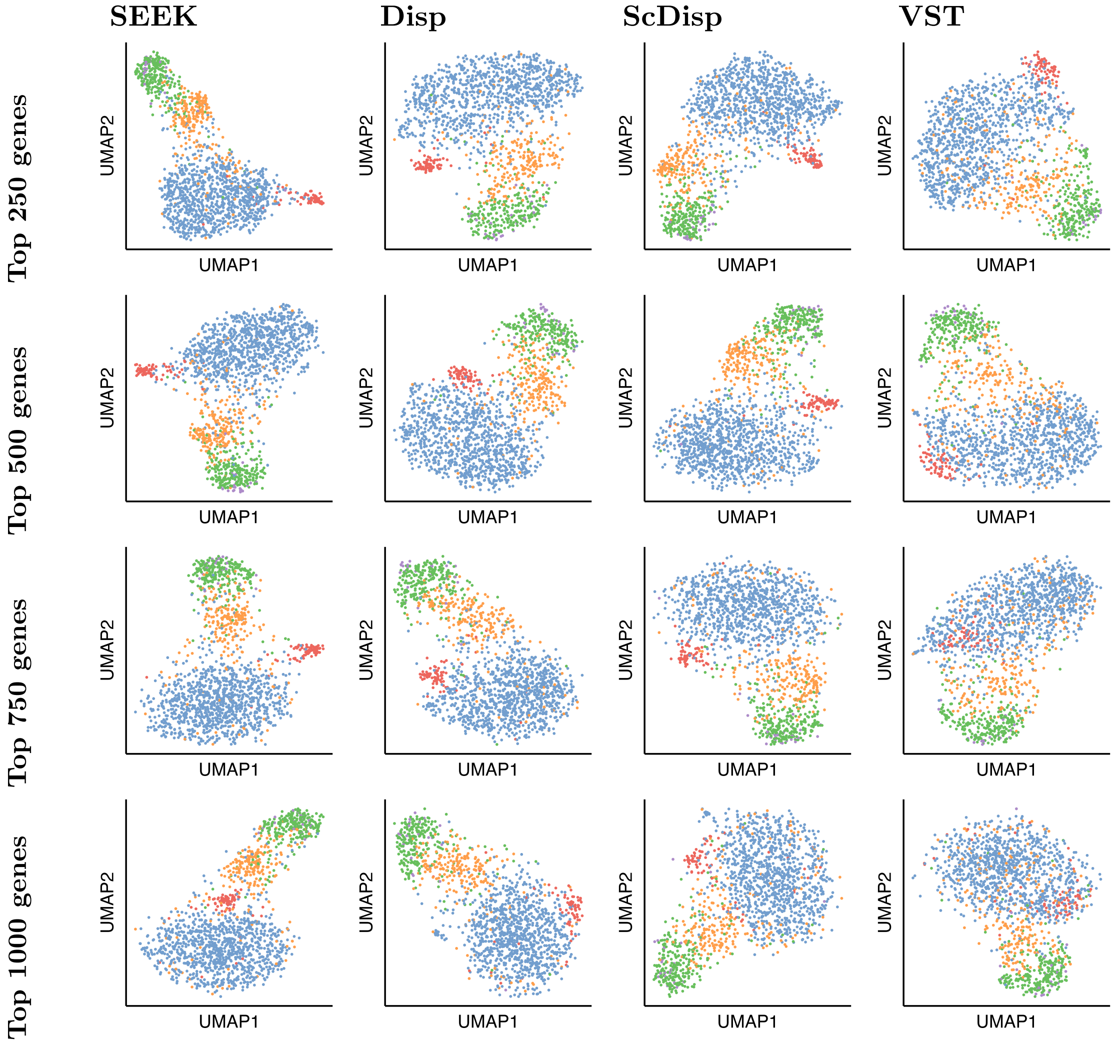


Supplemental Figure S28: UMAP embeddings of the thymus data with using the top $X$ genes. Columns correspond to the method used to select the top genes and rows correspond to the value of $X$.

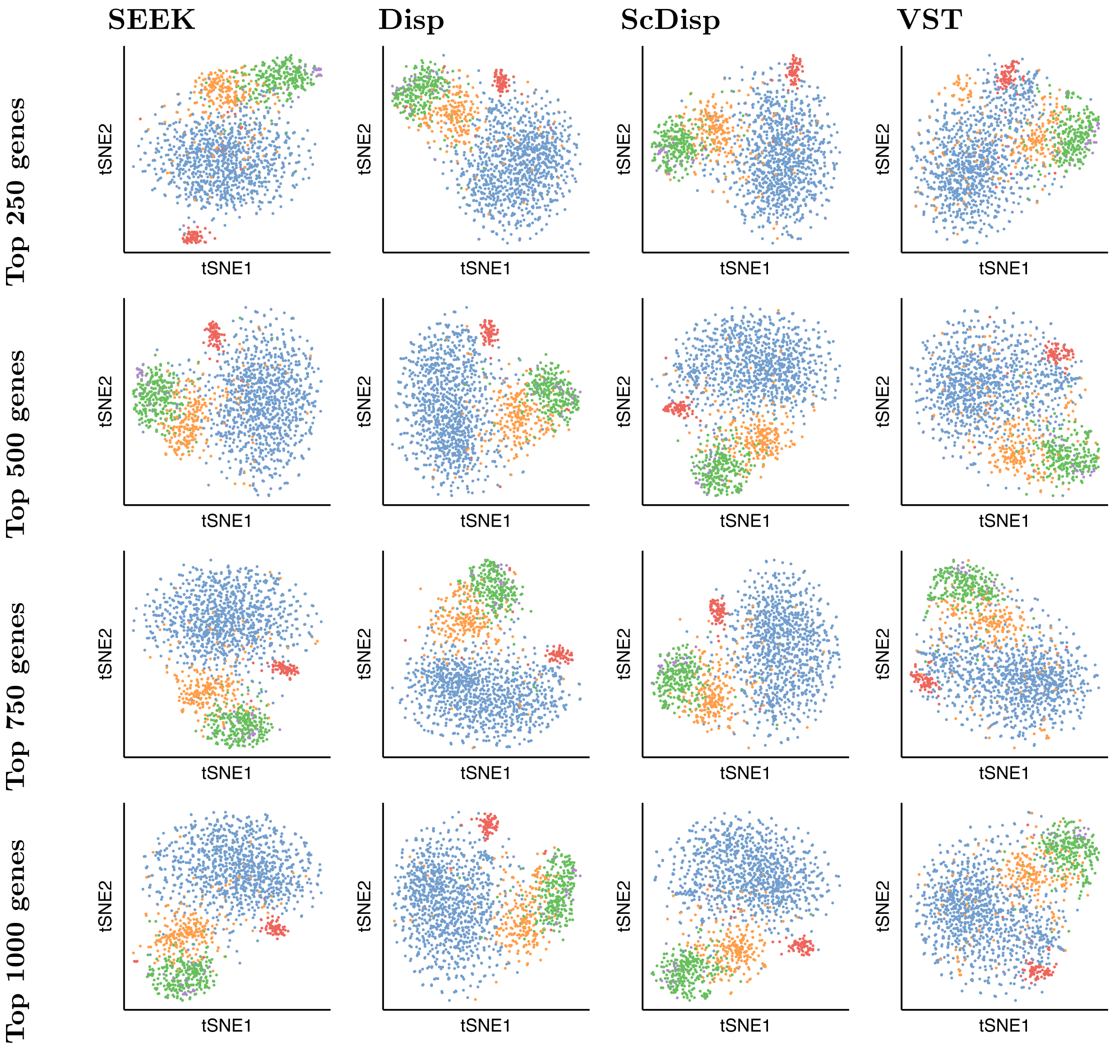


Supplemental Figure S29: t-SNE embeddings of the thymus data with using the top $X$ genes. Columns correspond to the method used to select the top genes and rows correspond to the value of $X$.

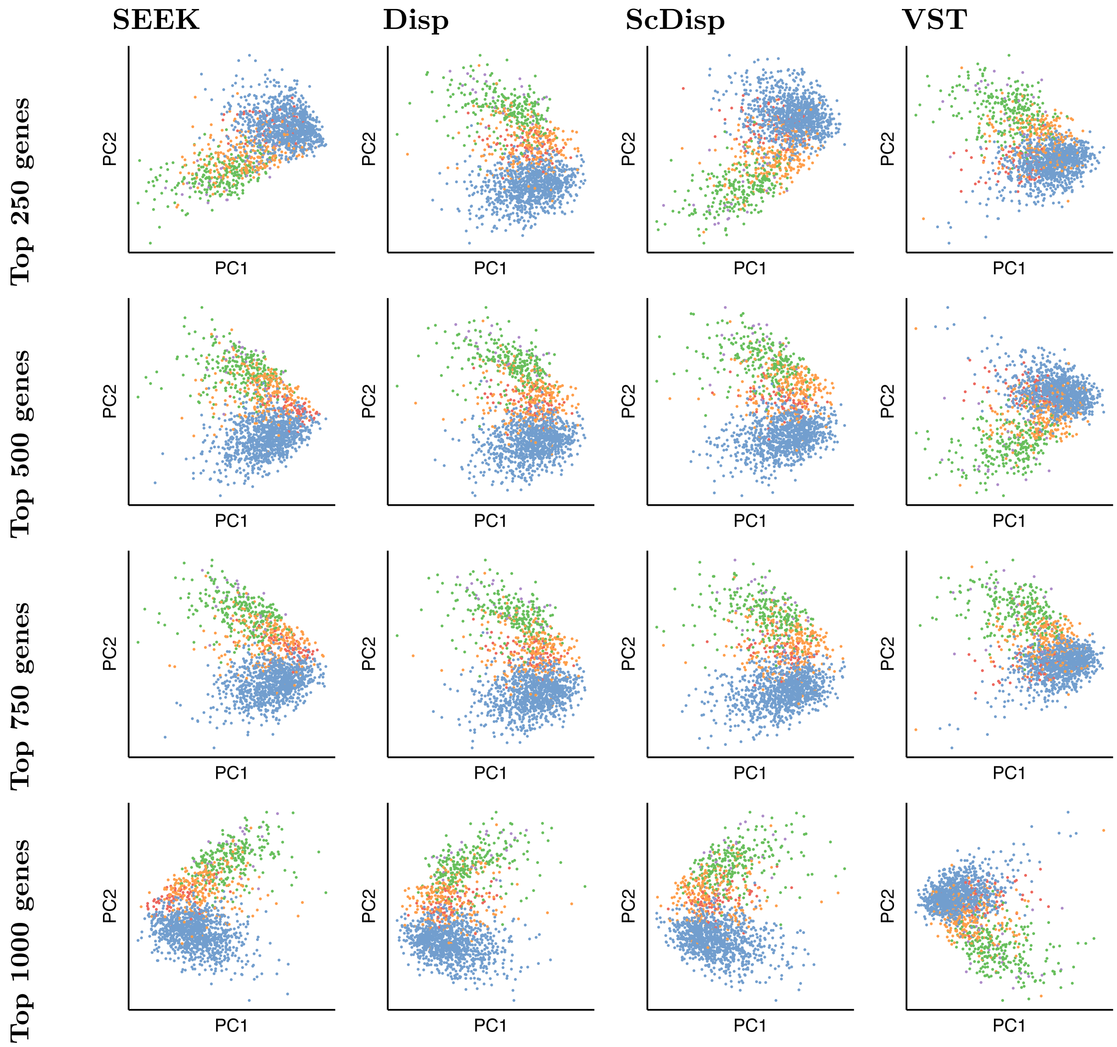


Supplemental Figure S30: PCA embeddings of the thymus data with using the top $X$ genes. Columns correspond to the method used to select the top genes and rows correspond to the value of $X$.
